# Single molecule long read sequencing resolves the detailed structure of complex satellite DNA loci in *Drosophila melanogaster*

**DOI:** 10.1101/054155

**Authors:** D. E. Khost, D. G. Eickbush, A. M. Larracuente

## Abstract

Satellite DNA (satDNA) repeats can make up a large fraction of eukaryotic genomes. These blocks of tandem repeats are rapidly evolving and have roles in genome stability and chromosome segregation. Their repetitive nature poses genome assembly challenges and has stymied progress on the detailed study of satDNA structure. Here we use single molecule real-time sequencing reads to assemble and study the genomic structure of two complex satDNA loci in *Drosophila melanogaster—260-bp* and *Responder*—with unprecedented resolution. We find that complex satDNAs are organized into large tandem arrays interrupted by transposable elements. The homogenized repeats in the array center suggest that gene conversion and unequal crossovers drive the concerted evolution of repeats, the degree to which differs among satDNA loci. Both satDNA arrays have a higher order organization that suggests recent structural rearrangements. These assemblies provide a platform for the evolutionary and functional genomics of satDNAs in pericentric heterochromatin.

## INTRODUCTION

Satellite DNAs (satDNAs; Kit 1961; Sueoka 1961; Szybalski 1968) are tandemly repeated DNAs frequently found in regions of low recombination (Charlesworth, et al. 1994; *e.g.* centromeres, telomeres, and Y chromosomes) that can make up a large fraction of eukaryotic genomes (Britten and Kohne 1968). SatDNA families are classified according to their repeat unit size and composition—simple satellites generally correspond to uniform clusters of small (*e.g* 1-10 bp) repeat units and complex satellites correspond to more variable clusters of larger (*e.g.* > 100 bp) repeat units. SatDNAs are highly dynamic over short evolutionary time scales (Lohe and Brutlag 1987b; Larracuente 2014). Changes in satDNA composition and abundance contribute to the evolution of genome structure (Charlesworth, et al. 1994), speciation (Yunis and Yasmineh 1971; Ferree and Barbash 2009) and meiotic drive (Henikoff, et al. 2001; Fishman and Saunders 2008). Early studies on satDNA (correctly) assumed that it must have some function in protecting against nondisjunction during chromosome segregation (Walker 1971) or a structural role in the nucleus (Yunis and Yasmineh 1971). However, subsequent studies suggested that satDNAs were inert “junk” (Ohno 1972) that expand in genomes due to selfish replication (Doolittle and Sapienza 1980; Orgel and Crick 1980; Orgel, et al. 1980). In the last 15 years, researchers across the fields of evolutionary, cell and molecular biology have accumulated evidence that some satDNAs have important functions (Dernburg, et al. 1996; Sun, et al. 1997; Csink and Henikoff 1998; Ferree and Barbash 2009; Hughes, et al. 2009; Zhu, et al. 2011; He, et al. 2012). However, the highly-repetitive nature of satDNA makes the detailed study of their loci difficult.

Gross-scale techniques such as density-gradient centrifugation and *in situ* hybridization demonstrate that satDNAs are organized into large contiguous blocks of repeats (Peacock, et al. 1974; Lohe and Brutlag 1986). Molecular assays based on restriction digest mapping indicate that satDNA blocks may be interrupted by smaller “islands” of more complex repeats such as transposable elements in *Drosophila* mini chromosomes (Le, et al. 1995; Sun, et al. 1997). While these methods have been useful in describing the overall structure of satDNA loci, detailed sequence-level analysis of these arrays is stymied by the shortcomings of traditional sequencing methods. Highly-repetitive arrays are unstable in BACs and cloning vectors (Brutlag, et al. 1977; Lohe and Brutlag 1986, 1987a)—in some cases they are even toxic to *E. coli* and thus are underrepresented in BAC libraries and among Sanger sequence reads (Hoskins, et al. 2002). Next generation short-read sequencing methods such as Illumina or 454 circumvent bacterial-based cloning related issues. These methods still pose a difficulty for repeat assembly because of PCR biases and short read lengths that result in the collapse of, or assembly gaps in, repetitive regions (Hoskins, et al. 2002; Schatz, et al. 2010). However, recent developments in single-molecule real-time (SMRT) sequencing (*e.g.* from Pacific Biosciences; PacBio; Eid, et al. 2009) address some of these shortcomings (Koren, et al. 2012; Chin, et al. 2013; Berlin, et al. 2015). With current sequencing chemistries, PacBio read lengths are ~16 kb on average but reach ~50kb, which can bridge repetitive regions not resolved with short read technology. While PacBio reads have a high error rate (~15%), because these errors are randomly distributed, several approaches can correct the reads for use in *de novo* assembly (Koren, et al. 2012; Ross, et al. 2013; Chaisson, et al. 2014; Lam, et al. 2014). Hybrid approaches use deep coverage from Illumina reads for error correction of the raw PacBio reads (Koren, et al. 2012). However, hybrid assemblies have difficulty dealing with regions that have large dips in short-read coverage, which can be caused by GC or sequence context biases known to affect Illumina data, resulting in breaks in the final assembly (Koren, et al. 2012; Chin, et al. 2013). More promising for the *de novo* assembly of repetitive regions are algorithms that use the PacBio reads themselves for self-correction (Koren, et al. 2012; Chin, et al. 2013). With sufficiently high read coverage (>50X), the longest subset of reads are corrected by overlapping the shorter reads and the corrected long reads are then used for contig assembly, which can produce assemblies that are more contiguous than hybrid assemblies (Berlin, et al. 2015; Chakraborty, et al. 2016). One popular package for PacBio assembly is the PBcR pipeline included in the Celera assembler. Earlier versions of the assembler (Celera 8.1) used a time-intensive all-by-all alignment step called BLASR to compute overlaps among the uncorrected reads, which accounts for >95% of runtime and is a significant bottleneck for larger genomes (Berlin, et al. 2015). More recent versions (Celera 8.2+) use the newly developed MinHash Alignment Process (MHAP) algorithm to overlap and correct the reads, which is several orders of magnitude faster than BLASR (Berlin, et al. 2015). Recently, the Celera assembler was forked to create the Canu assembler, which specializes in assembling PacBio error-prone reads (Berlin, et al. 2015; https://github.com/marbl/canu). Similar to the Celera 8.2+ pipeline, Canu uses the MHAP algorithm for fast read alignment and assembly.

Assembly quality is most often evaluated based on increased overall contiguity and ability to close gaps in the euchromatin. Here, we assess the utility of long-read SMRT sequencing approaches for the accurate assembly of historically challenging repetitive regions near the centromere. We experiment with MHAP, Canu and BLASR-based PacBio *de novo* assembly methods to assemble satDNA regions in the pericentric heterochromatin of the *D. melanogaster* genome. We focus on two complex satDNA loci— *Responder (Rsp)* and *260-bp—* and assess assembly quality through molecular and computational validation. *Rsp* is a satDNA that primarily exists as a dimer of two related 120-bp repeats, referred to as *Left* and *Right*, on chromosome *2R* (Wu, et al. 1988; Pimpinelli and Dimitri 1989; Houtchens and Lyttle 2003; Larracuente 2014). *Rsp* is well-known for being a target of the selfish male meiotic drive system *Segregation Distorter* (reviewed in Larracuente and Presgraves 2012). *260-bp* is a member of the *1.688* family of satellites and is located on chromosome *2L* (Abad, et al. 2000). Using high-coverage (~90X) PacBio data for *D. melanogaster*, we determine the optimal assembly protocols for complex satDNA loci and provide a detailed, base pair-level analysis of the *Rsp* and *260-bp* complex satDNAs.

## RESULTS

### Rsp and 1.688 FISH

To confirm the gross-scale genomic distribution of *Rsp* and *260-bp* satellites in the sequenced strain (ISO1), we performed multi-color fluorescence *in situ* hybridization (FISH) on mitotic chromosomes. *Rsp* is located in the pericentric heterochromatin on chromosome *2R* (Fig 1; S1), proximal to clusters of *Bari-1* repeats (Fig S1), in agreement with previous studies (Caizzi, et al. 1993) and the PacBio assemblies. The *260-bp* satellite is in *2L* heterochromatin (Fig 1). The *260-bp* probe cross-hybridizes with other members of the *1.688* family: *353-bp* and *356-bp* on chromosome *3L* as well as *359-bp* on chromosome *X* (Fig 1).

**Figure 1:**
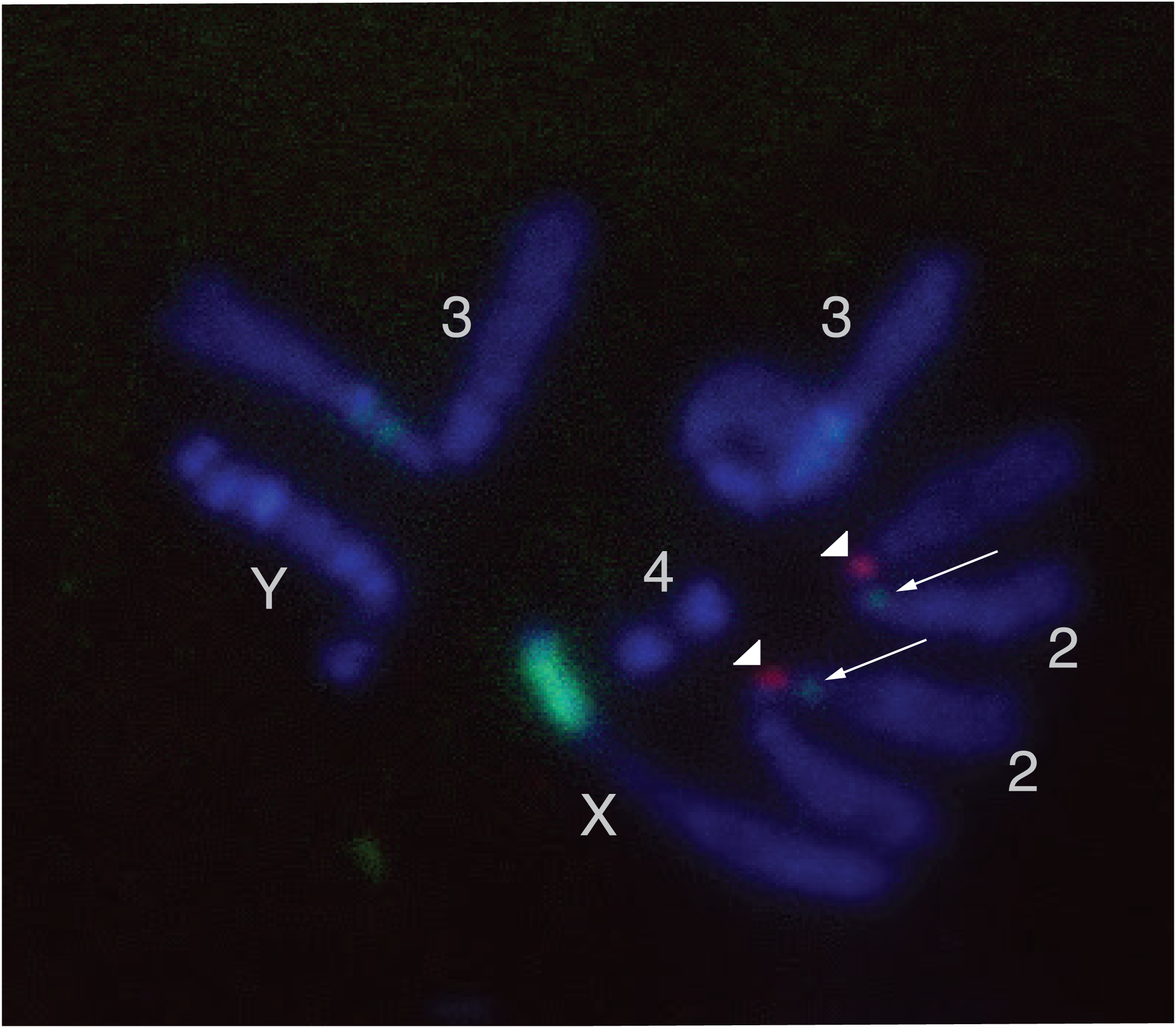
FISH image of *D. melanogaster* mitotic chromosomes showing *Rsp* and *260-bp* satDNAs. DNA is stained with DAPI (blue), *Rsp* is indicated by an avidin-rhodamine probe (red; arrowhead) and *260-bp* by an anti-digoxigenin probe (green; arrow). The *260-bp* probe also targets other members of the *1.688* satellite family on the X and 3L chromosome arms.

### Optimal approaches to complex satellite DNA assembly

Our goal was to determine the best methods for assembling arrays of complex satellites. We compared *de novo* PacBio assemblies generated using different methods and parameters (both our own and existing assemblies) and evaluated them based on the contiguity of complex satellite sequences. We generated *de novo* PacBio-only assemblies using the Celera 8.3 and 8.2 PBcR pipelines (referred to as “MHAP”) using a range of parameters (Table S1). We generated assemblies with the experimental FALCON diploid assembler that yielded highly fragmented assemblies that we will not discuss further (Table S2). We also experimented with the recently developed Canu 1.2 assembler using a range of error rates for the overlapping steps (referred to as “Canu” followed by the error rate; Table S3). Lastly, to determine which step is most important to proper assembly, we also generated two assemblies using the Canu and Celera 8.3 assemblers but with pre-corrected reads from the computationally-intensive BLASR method (referred to as Canu-corr and BLASR-corr Cel8.3, respectively). For the *260-bp* locus, all MHAP and Canu assemblies that we built using the diploid/large genome parameters, as well as the PBcR-BLASR assembly, recovered a 1.3-Mb contig that contains ~230 *260-bp* repeats spanning ~75 kb (Table 1). The other contigs containing *260-bp* have < 10 copies, or are short contigs made up of only satellite sequence. Our MHAP assemblies tended to produce these short contigs comprised entirely of *Rsp* or *1.688* family satellites, which were not present in the Canu, PBcR-BLASR and the BLASR-corr Cel8.3 assemblies. In contrast to the *260-bp locus*, the *Rsp* locus on *2R* was more variable between the different assembly methods. MHAP assemblies that lacked the diploid/large genome parameters and Canu assemblies with more stringent error rates produced a fragmentary locus consisting of several contigs with ~200-300 *Rsp* repeats per contig. The PBcR-BLASR and BLASR-corr cel8.3 assemblies each contained a single contig with ~1000 *Rsp* repeats, and whose distal end matched the *Rsp* locus in the latest release of the *D. melanogaster* reference genome (Release 6.03, which contains only ~300 copies). Our genomic Southern results are consistent with this number while our pulse field gel analysis suggested a somewhat higher number of repeats. The latter could be due to the different conditions under which the gels were run (Fig S2). Our slot blot analysis estimating the relative abundance of *Rsp* in ISO1 compared to three genotypes with previously published estimates of *Rsp* copy number (*cn bw, lt pk cn bw* and *SD*; Wu, et al. 1988; Supplemental methods) is also consistent with our assembly. It is important to note that while relative abundances are accurately estimated, we believe that the precise quantification of *Rsp* copy number using any hybridization-based method is not feasible due to sequence variability in the repeats at the *Rsp* locus (*e.g.* see Houtchens and Lyttle 2003). *Rsp-* containing BACs mapping to *2R* heterochromatin align with >99% homology to the distal portion of the locus. Several of our MHAP assembly parameter combinations (*e.g*. MHAP 20_1500_25X) and Canu assemblies with a more permissive error rate (*e.g.* Canu 4%) also produced a *Rsp* locus with ~1000 repeats, similar to PBcR-BLASR and BLASR-corr cel8.3 (Table 1). Notably, the Canu 4% assembly resulted in a contig with ~1100 *Rsp* repeats but also extending another ~250 kb distal to the *Bari1* repeats (total contig length ~587 kb, Fig. S3). However, while the total locus size and number of repeats were roughly consistent between the PBcR-BLASR, BLASR-corr Cel 8.3, Canu and MHAP assembly methods, close examination revealed rearrangements in the central *Rsp* repeats between these assemblies. Specifically, the *Rsp* locus in the Canu-corr and BLASR-corr Cel 8.3 assemblies had an inversion relative to the PBcR-BLASR assembly, as did several of our MHAP assemblies. There were also various medium-to-long indels over the center of the locus between the different assemblies.

**Table 1:**
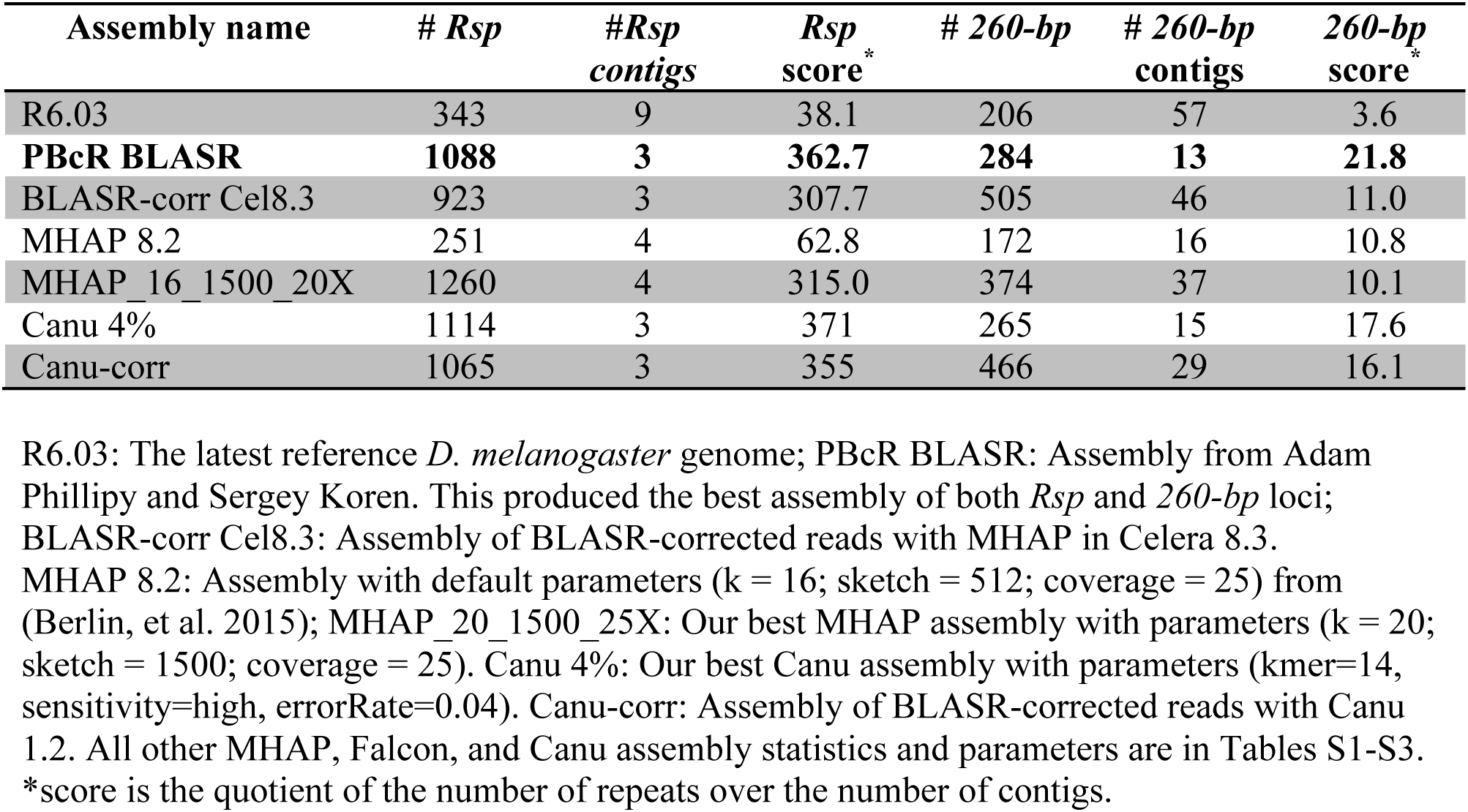
Summary of *Rsp* and *260-bp* repeats counts for a subset of assemblies. Counts are for all assembled repeats in any genomic contig.

| Assembly name | # *Rsp* | #*Rsp*<br>contigs | *Rsp*<br>score<sup>*</sup> | # 260-bp | # 260-bp<br>contigs | 260-bp<br>score<sup>*</sup> |
| --- | --- | --- | --- | --- | --- | --- |
| R6.03 | 343 | 9 | 38.1 | 206 | 57 | 3.6 |
| **PBcR BLASR** | **1088** | **3** | **362.7** | **284** | **13** | **21.8** |
| BLASR-corr Cel8.3 | 923 | 3 | 307.7 | 505 | 46 | 11.0 |
| MHAP 8.2 | 251 | 4 | 62.8 | 172 | 16 | 10.8 |
| MHAP_16_1500_20X | 1260 | 4 | 315.0 | 374 | 37 | 10.1 |
| Canu 4% | 1114 | 3 | 371 | 265 | 15 | 17.6 |
| Canu-corr | 1065 | 3 | 355 | 466 | 29 | 16.1 |

R6.03: The latest reference *D. melanogaster* genome; PBcR BLASR: Assembly from Adam Phillippy and Sergey Koren. This produced the best assembly of both *Rsp* and 260-bp loci; BLASR-corr Cel8.3: Assembly of BLASR-corrected reads with MHAP in Celera 8.3. MHAP 8.2: Assembly with default parameters (k = 16; sketch = 512; coverage = 25) from (Berlin, et al. 2015); MHAP\_20\_1500\_25X: Our best MHAP assembly with parameters (k = 20; sketch = 1500; coverage = 25). Canu 4%: Our best Canu assembly with parameters (kmer=14, sensitivity=high, errorRate=0.04). Canu-corr: Assembly of BLASR-corrected reads with Canu 1.2. All other MHAP, Falcon, and Canu assembly statistics and parameters are in Tables S1-S3. \*score is the quotient of the number of repeats over the number of contigs.

### Molecular and computational validation of the Rsp locus

To distinguish between the possible configurations of the locus, we mapped high coverage Illumina and raw PacBio reads to the assemblies containing ~1000 *Rsp* copies (PBcR-BLASR, BLASR-corr Cel 8.3, Canu 4%, and our example MHAP assembly 20_1500_25X). Each MHAP assembly, including the ones with highest contiguity of *Rsp*, had dips in coverage across the *Rsp* locus, suggesting that they might be mis-assembled (*e.g.* Fig S4). Similarly for the Canu 4% assembly, both the PacBio and Illumina mapped reads have sharp dips in coverage near the center of the *Rsp* locus (Fig S5). Several regions that have zero Illumina read coverage also have very low PacBio coverage (<10 reads). In contrast, the PBcR-BLASR and BLASR-corr cel8.3 assemblies had uniform coverage across the contig for both the Illumina and PacBio reads (*e.g.* Fig S6-S8). We therefore focus on these two assemblies. While both were well supported by read mapping, aligning the PBcR-BLASR assembly against the BLASR-corr cel8.3 assembly showed an inversion in the central segment of the major *Rsp* locus (Fig S9). To determine the correct orientation, we designed long PCR primers that should amplify a 15kb product based on the PBcR-BLASR assembly and no product based on the BLASR-corr Cel8.3 assembly (indicated in Fig 2A; primer pair 3). We obtained a 15kb fragment, which we excised and digested with several restriction enzymes; Southern analysis of these digests was as predicted from the PBcR-BLASR assembly (Fig S2A). In addition, we performed Southern blot analysis of restriction enzyme digested genomic DNA to look at large segments across the entire major *Rsp* locus; these results also supported the PBcR-BLASR assembly (Fig S2B-C). While the predicted restriction digest patterns for the major *Rsp* locus are also consistent with the most proximal and distal regions in Canu 4% assembly, we found large dips in PacBio and Illumina read coverage over the center of the locus in this assembly. For the major *Rsp* locus, the time-intensive BLASR correction step appears to be required, and we therefore use the PBcR-BLASR assembly for subsequent analysis.

**Figure 2:**
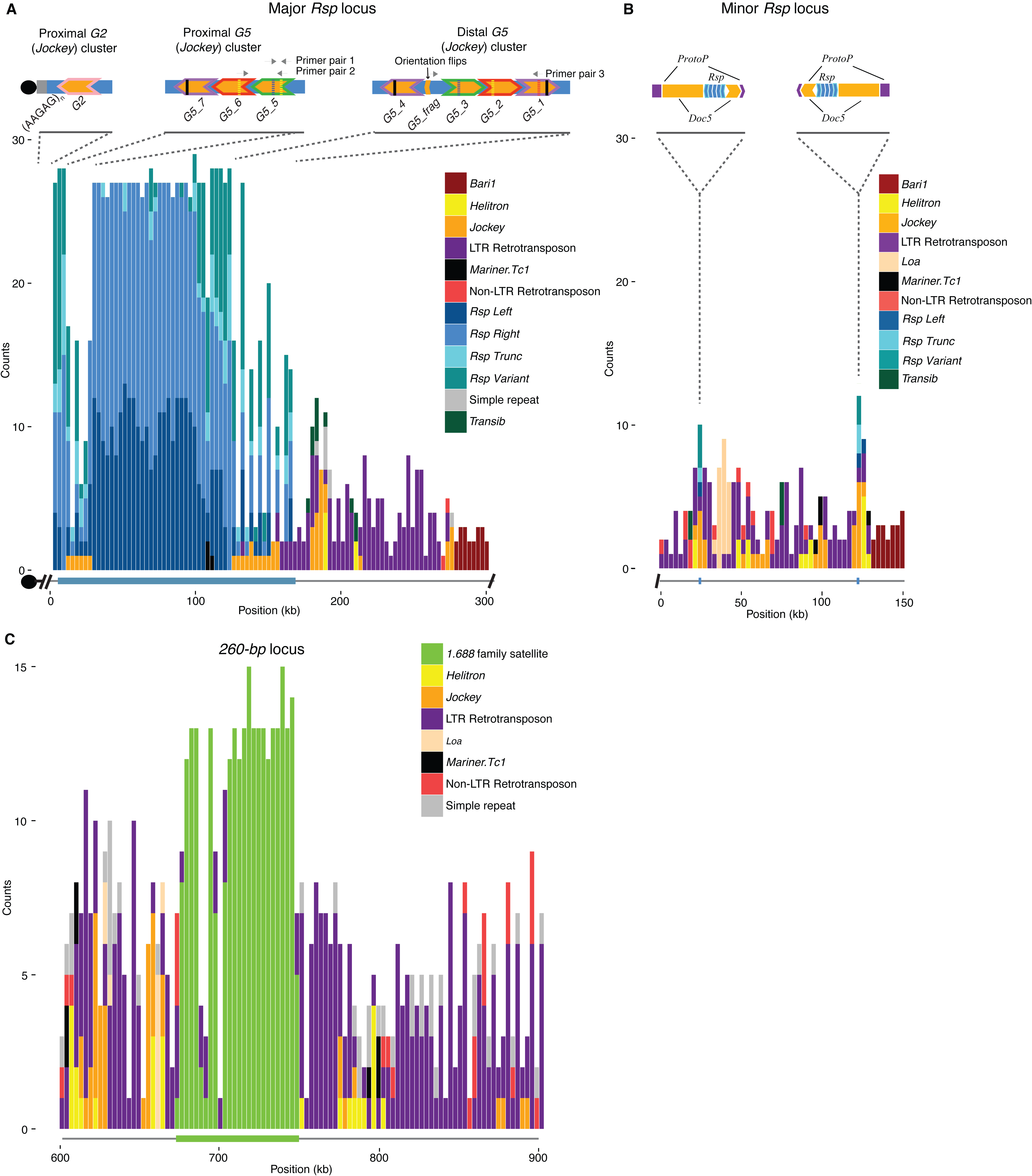
Maps of complex satDNAs contigs. Counts for each repetitive element family in our custom Repbase library were plotted in 3-kb windows across each contig. **A)** *Rsp* locus on chromosome *2R*. Blue bars correspond to *Rsp Left, Right* or *Variant* or *Truncated* repeats, while other colors correspond to various TE families as indicated to the right of each contig. *Rsp* spans ~170 kb of the 300-kb contig (thick blue line below x-axis). Above the plot is a schematic showing the orientation of two *G5* clusters flanking the *Rsp* locus and a separate contig containing *Rsp* and the Jockey element *G2*, which is directly adjacent to centromeric AAGAG repeats. The colors of the chevron outlines indicate the *G5* elements with the highest degree of identity with one another. Solid and dashed lines within the insertions show the approximate locations of shared insertions or deletions (respectively). Several configurations of indels are unique, such as the two in *G5_5* or the deletion in *G5_1*, which allows verification of the cluster. The *G2* contig links the *Rsp* locus to what appears to be the chromosome *2* centromere (Black circles). **B)** Minor *Rsp* locus on *2R*. The inset shows the detailed orientation of the two clusters (5 *Rsp* repeats per cluster, ~100 kb apart); direction of arrows indicated the relative orientation of the elements. The *Rsp* repeats (blue chevrons) are nested within *Doc5* (orange chevrons) insertions, which are in turn nested within insertions of a transposon known as *ProtoP* (purple chevrons). The clusters of *Rsp+Doc5* share approximately 96% sequence identity with one another, and are in an inverted orientation. **C**) *260-bp* locus on chromosome *2L*. Only the area surrounding the *260-bp* array is shown (300 kb of ~1.1-Mb contig). The *260-bp* locus spans ~70 kb of the 1.1-Mb contig (green line below x-axis) and is interrupted with *Copia* elements.

To evaluate if there is bias in the error rate over our satDNA loci, we compared the rate of single nucleotide substitutions and indels over the contigs containing *Rsp* and *260-bp* vs the rest of the assembly using Pilon with high-coverage Illumina data. The nucleotide substitution error rate for both satDNA loci are close to the median of the empirical cumulative distribution function (ecdf) of the rate for all contigs (Fig S10). The indel error rates are in the first quartile of the ecdf, but still not significantly different (Fig S11).

### Structure of Rsp and 260-bp loci

We find that a single 300-kb contig contains most of the major *Rsp* locus, while a 150-kb contig contains a minor locus directly distal to the major locus. The minor locus contains *Bari1* repeats and two small clusters of variant *Rsp* repeats (five repeats per cluster, ten total) separated by ~100kb (Fig 2B). In the Canu 4% assembly both the major and minor *Rsp* loci are contained within a single contig (Fig S3). Interestingly, these small *Rsp* clusters are each inserted in the middle of a *Doc5* transposon (which is itself inserted in a *ProtoP* element). This *Rsp-Doc5-ProtoP* feature is duplicated in inverted orientation ~100 kb away (Fig 2B) and ~96% identical to one another.

The major *Rsp* locus is ~170 kb and contains ~1050 *Rsp* repeats, which are interrupted by transposable element sequences at the centromere proximal (left) and distal (right) ends of the array. The presence of the *Bari1* repeats at the distal end of the contig agrees with our FISH analysis (Fig S1) and previous studies (Wu, et al. 1988; Caizzi, et al. 1993). However, because the proximal end of the contig terminates in *Rsp*, it may be missing the most centromere-proximal repeats. We found seven raw uncorrected PacBio reads containing large (up to 6-kb) blocks of both tandem *Rsp* repeats and the centromeric AAGAG simple satellite, confirming the presence of additional repeats and suggesting that the major *Rsp* locus is directly centromere-adjacent (Lohe, et al. 1993). The AAGAG+*Rsp* reads were not present in the error-corrected PacBio reads, and due to the high error rate of the uncorrected reads, we could not compare the AAGAG-adjacent *Rsp* repeats to our contig. However, the AAGAG+*Rsp* reads also contain a single *Jockey* element insertion called *G2*, which we used to identify 11 error-corrected reads containing *Rsp* and the *G2* insertion that link these most centromere-proximal repeats to the rest of the locus (Fig 2A). We created a contig from the 11 error-corrected reads (Fig S12) that, when combined with the AAGAG+*Rsp* raw reads, suggests that our 300-kb contig is missing ~22 kb of sequence containing ~200 *Rsp* repeats. These *Rsp* elements are most similar to the proximal-most repeats in our 300-kb contig (Fig 3A), providing additional support that they indeed correspond to the centromere-proximal repeats.

**Figure 3:**
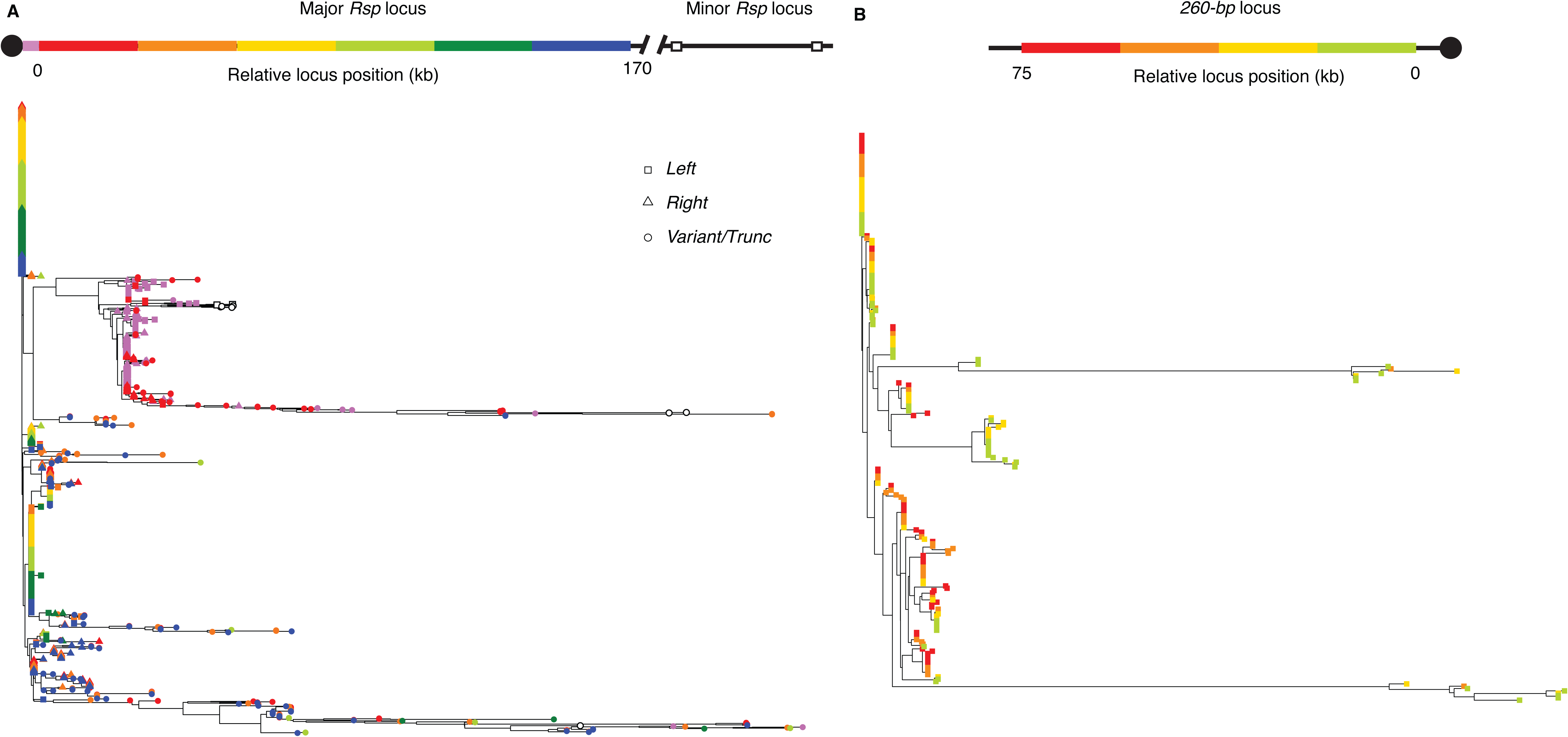
Neighbor-joining tree of complex satDNA monomers. **A.)** *Rsp* repeats in the chromosome *2R* locus. Repeats were divided into bins where each bin contains 1/6^th^ of the locus, or ~180 repeats/bin. Tip color corresponds to position in the array (red is most centromere proximal, blue is most centromere distal). The tip symbol indicates if the repeat is *Left* (square)*, Right* (triangle), or *Variant/truncated* (circle). Repeats corresponding to the *G2* contig containing the most centromere-proximal repeats are indicated in pink. Note that these repeats cluster with the repeats on the proximal end of the *Rsp* contig (red), supporting their location adjacent to the centromere. The minor *Rsp* locus is show in white, and clusters with the centromere-proximal *Rsp* repeats from the major locus. **B.)** *260-bp* repeats in the chromosome *2L* locus. Repeats were divided into bins where each bin contains 1/4^th^ of the locus, or ~57 repeats/bin. Tip color corresponds to position in the array (red is most centromere proximal and green is most centromere distal).

Satellites tend to undergo concerted evolution—unequal exchange and gene conversion homogenize repeat sequences within arrays (Dover 1982; Charlesworth, et al. 1994; Dover 1994). To test the hypothesis that *Rsp* undergoes concerted evolution, we examined the relationship between genetic and physical distance within the *2R* array. We built neighbor-joining trees for each satellite family using each full-length repeat monomer (Fig 3). We find a pattern consistent with concerted evolution: two large clades of nearly identical repeats corresponding to the *Right* and *Left Rsp* repeats consist mainly of repeats from the center of the array. In contrast, the *Variant* repeats have longer branch lengths and tend to occur toward the proximal and distal ends of the array (Fig 3A). To examine the higher-order structure of the array, we studied the distribution of all unique repeats sequences across the locus according to their abundance (Fig 4A). The ~1050 *Rsp* repeats on the main contig correspond to ~480 unique variants. Consistent with our phylogenetic analysis, low copy number *Rsp* repeats tend to dominate the ends of the array, while higher copy number variants dominate the center of the array (Fig 4A).

**Figure 4:**
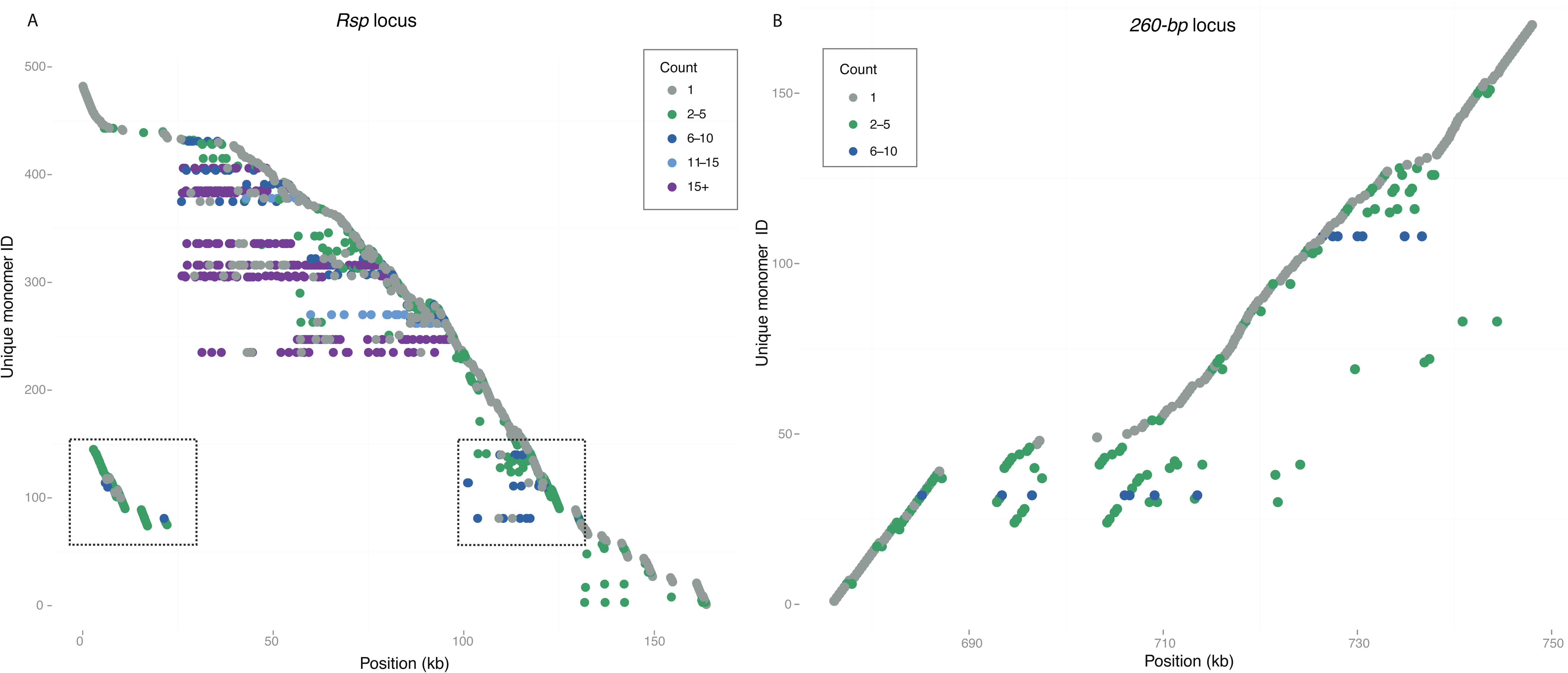
Distribution of satDNA sequence variants across loci. Each row corresponds to a unique monomer, while the x-axis shows the position of that monomer sequence in the array. The color of the point indicates the copy number of each monomer in the array. **A)** The *Rsp* locus on *2R*. Several high copy number *Rsp* variants dominate the center of the array (purple and blue), with the low frequency and unique sequences found more towards the proximal and distal ends (gray and green). One cluster of repeats is duplicated on either side of the array (boxed). **B.)** The *260-bp* locus on *2L*. The majority of repeats occur only once, while a few variants have intermediate copy number.

There are several TE insertions within the major *Rsp* array located towards the proximal and distal ends of the locus. The homogenized *Rsp* repeats in the center of the array are flanked by two nearly identical clusters of *G5 Jockey* elements (Fig 2A). These *G5* repeats form their own clade with respect to the other *G5* insertions in the genome and have a high degree of similarity to one another (Fig S13). They have a complicated orientation, with each repeat having a near 99% identical (albeit inverted) match on the opposite side of the locus ~100 kb away (Fig 2A). Despite the similarity between the two clusters, there are several unique configurations of indels in each that distinguish them. We examined the pileup of raw PacBio reads over sets of long indels found in the *G5* clusters and identified 8 and 20 individual long reads that spanned the unique configuration of indels in the proximal *G5* cluster (*G5-5* and *G5-6*, Fig 2A) and distal *G5* cluster (*G5-3* and *G5-2*, Fig 2A), respectively. This suggests that the proximal cluster actually exists and is not an error in the assembly of the distal cluster. For further confirmation, we designed PCR primers complimentary to the unique indels in the proximal cluster (Fig 2A), which return products of the expected size (data not shown). The *Rsp* elements surrounding the *G5* elements also show a mirrored structure (Fig 4A). Interestingly, one 1.7-kb stretch of inter-*G5 Rsp* repeats is repeated three times, which suggests a complex series of duplication and inversion within the *G5* cluster. The *Rsp* repeats are oriented on the same strand across most of the array, but they flip orientation at the fragmentary *G5* element, mirroring what we see with *G5* elements (Fig 2A). Thus the inversion did not occur only in the local area around the *G5*s, but across the entire proximal end of the contig.

The *260-bp* locus on chromosome *2L* is fully contained within a 1.2-Mb contig and contains 230 repeats interrupted by identical *Copia* transposable elements (Fig 2B). Unlike *Rsp*, the *260-bp* satellite array lacks the homogenized center and has more variant sequences (Fig 3B). The *260-bp* satellite has relatively more unique variants than *Rsp*: the 230 monomers correspond to 170 unique variants, and there are fewer high copy number variants (Fig 4B).

## DISCUSSION

### Assembly methods for complex satellites

For large complex centromeric repeats, such as human centromeres, the complete assembly of a contiguous stretch of repeats has not been possible with current technologies (Miga 2015). Instead, human centromere composition can be inferred using clever graph-based modeling strategies (Miga, et al. 2014). In contrast, single molecule sequencing has produced assemblies of more tractable, but still challenging highly repetitive genomic regions (Chaisson, et al. 2014; Carvalho, et al. 2015; Krsticevic, et al. 2015), including some plant centromeres (VanBuren, et al. 2015; Wolfgruber, et al. 2016). However, validation of these assemblies is difficult. Here, we annotate accurate *de novo* assemblies of two complex satDNAs in *D. melanogaster* using SMRT PacBio sequencing reads, allowing us to examine the detailed spatial distribution of elements within these arrays for the first time. We found that assemblers differed in their ability to produce a complete assembly for the two satellites: while the *260-bp* locus assembly was consistent between almost all PacBio assembly methods, the larger *Rsp* locus required the time-intensive BLASR correction algorithm for an accurate assembly. We validated the major features of this PBcR-BLASR *Rsp* assembly through extensive molecular and computational approaches and, with some manual scaffolding, were able to extend the assembly to what may be the junction between *Rsp* and the chromosome *2* centromeric AAGAG satDNA. There are four features of the *Rsp* locus that could present a particular challenge for *de novo* assembly, especially for MHAP- and FALCON-based methods: *1)* it is large (more than twice the size of the *260-bp* locus); *2)* it appears to be centromere-adjacent (Pimpinelli and Dimitri 1989), with AAGAG repeats directly proximal to the *Rsp* cluster; *3)* the array center is occupied by a contiguous stretch of nearly identical repeat variants; and *4)* these repeats are flanked by nearly identical TEs in inverted orientation. In contrast, the *260-bp* locus is smaller, lacks large runs of totally identical sequence, and does not have as complex higher-order organization. In addition to struggling with the major *Rsp* satDNA locus, we found that even our most contiguous MHAP assemblies produced short contigs consisting entirely of what we believe are extraneous repeats. Despite these caveats, we recover the gross-scale organization of the locus with our best MHAP and Canu parameter combinations. We found that the MHAP assembler requires the “diploid/large genome” parameters to produce a contiguous major *Rsp* locus. These parameters allow for a great fraction of errors in the overlapping steps during error-correction, consensus calling, and unitig construction/assembly. Similarly, we found that a more strict (lower) error rate with the Canu resulted in more fractured satDNA loci, while allowing a more lenient (higher) error rate produced more contiguous loci, but at the cost of increased assembly time. In addition, the Canu 4% error rate assembly produced a major *Rsp* locus in an orientation matching the validated PBcR-BLASR locus. However, read mapping indicates that the center of the locus is misassembled. Notably, areas with no support from the Illumina reads have very low PacBio coverage, suggesting that variant or error-prone reads can create spurious overlaps in ambiguous regions. Thus, the MHAP and Canu assemblies show that a more permissive error threshold allows for better assembly of complex satDNA loci, but there is a trade-off with accuracy. The faster MHAP and Canu approaches may offer a reasonable starting point for determining the structure of difficult repetitive loci.

One explanation for why BLASR generates superior assemblies of certain satDNA loci as compared to MHAP and Canu is the difference in how the two methods calculate overlaps. The MHAP algorithm converts the k-mers for each read into an integer “fingerprint,” which are collected into a set (sketch) representing the whole sequence that can be easily compared to other reads to generate an overlap; in other words, it does not actually perform an alignment (Berlin, et al. 2015). BLASR, in contrast, generates a computationally expensive but sensitive all-by-all alignment of all the reads, which may imply full alignment of the reads is necessary for these regions. The fractured assemblies generated by MHAP suggest that, even when using large hash sizes, this method not sensitive enough for complex satDNAs. This finding is in contrast with recent work evaluating the repeat-rich Mst77Y region on the Drosophila Y chromosome, which found that the region is mis-assembled in the PBcR assembly but correct in the MHAP (Krsticevic, et al. 2015). Another study showed hybrid PacBio assembly combined with MHAP assembly produced the most contiguous assembly for *D. melanogaster* (Chakraborty, et al. 2016). Thus optimal PacBio assembly methods seem to be dependent on the region analyzed, and careful, independent verification of the assembly is important. We find that slower but more sensitive overlapping is required for base pair-level resolution of large complex satDNA loci like *Rsp*, while MHAP and Canu are sufficient for smaller, less homogeneous complex satDNA loci (such as *260-bp*). While the latest reference genome (release 6 assembly; Hoskins, et al. 2015) offered an impressive improvement in the assembly of pericentric regions over previous releases, the *de novo* PacBio assembly methods evaluated here (MHAP, Canu and PBcR-BLASR) produced more complete and contiguous assemblies of these complex satDNAs. No assembly method allowed us to resolve centric heterochromatin, which is enriched for simple satellite sequences. However, we did detect low coverage junctions between *Rsp* and the adjacent AAGAG repeats that occupy the centromere of chromosome *2*. We find a general reduced representation of simple satellite-rich raw reads, making it difficult to extend our assembly into the centromere. This apparent bias against raw reads derived from simple repeats has two potential explanations: *1)* PacBio sequencing is subject to a bias that is difficult to measure because it occurs in the most highly repetitive regions of the genome; and/or *2)* the inherent structural properties of some highly repetitive DNAs subject these sequences to misrepresentation in library preparation (*e.g.* non-random chromosome breakage during DNA isolation or library preparation). Therefore, the assembly of some simple tandem repeats still pose a significant challenge for PacBio-based assembly methods.

### Structure of complex satDNA loci

Consistent with gross-scale structural analyses of satellite DNA (Brutlag, et al. 1977; Carlson and Brutlag 1977; Lohe and Brutlag 1987a; Lohe, et al. 1993; Le, et al. 1995; Sun, et al. 1997), we find that *Rsp* and *260-bp* have uninterrupted blocks of homogeneous repeats alternating with “islands” of complex DNA. For both of these complex satDNAs, TE insertions cluster together towards the array ends. The TEs in and around the locus tend to be full-length and similar to euchromatic copies, suggesting recent insertion. What gives rise to this structure? Repetitive tandem arrays are thought to expand and contract *via* unequal crossing over (Smith 1976), which along with gene conversion can homogenize the array and lead to a pattern of concerted evolution (Dover 1982; Charlesworth, et al. 1994; Dover 1994). The localization of the TEs in islands near the proximal and distal ends of the locus is consistent with the “accretion model”, which predicts that repeated unequal exchange over the array should push TEs together and towards the ends of an array (McAllister and Werren 1999). The organization of the sequence variants across the locus and the degree of homogeneity differs between *Rsp* and *260-bp*. The center of the *Rsp* locus is highly homogeneous and dominated by a few high-copy number variants, while the *260-bp* locus is comprised mostly of low-copy number or unique repeats. These contrasts may simply be due to a difference in size of the two satDNAs, or more recent unequal exchange and gene conversion at the major *Rsp* locus. As exchange breakpoints are more likely to occur within repeats rather than perfectly at the junction between two repeats, the lack of truncated repeats within the array center suggests that any unequal exchange event involved a large chunk of the array. The nearly-identical *G5* elements flanking the major *Rsp* array suggest a complicated rearrangement, likely involving duplication and an inversion. The high degree of similarity between the clusters is unlikely a result of gene conversion—the clusters are ~100 kb apart and studies in rice have shown that rates of gene conversion decrease as a function of distance between elements (Xu, et al. 2008). Instead, the intervening *Rsp* locus may have recently expanded. Interestingly, we see a similar structure at the minor *Rsp* locus directly distal to the main locus: two small clusters of repeats are located 100 kb apart in an inverted orientation. Both clusters have 5 *Rsp* repeats inserted in the middle of *Doc5 Jockey* elements, which themselves interrupt *ProtoP* elements. In each case, the *Doc5* and *Rsp* elements are inserted in the same site, making it unlikely that the insertions occurred independently; instead, the entire *Rsp-Doc5-ProtoP* unit duplicated and inverted and are now separated by 100 kb. Because the *Rsp* repeats clearly interrupt the *Doc5* elements, it does not appear that the movement of *Rsp* was mediated by TE activity. We speculate that intra-sister chromatid exchange events in the major *Rsp* locus—in this case, the most centromere-proximal repeats (Fig 3A)—may have generated an extrachromosomal circular DNA, perhaps amplified through rolling circle replication, and re-integrated distal of the main cluster, in the middle of *Doc5.* Analogous events may seed the movement and expansion of satDNAs to new genomic regions. D*e novo* PacBio assembly methods allow for exciting progress in studying the structure of previously inaccessible regions of the genome in unprecedented detail. We show here that some complex satDNA loci are tractable models for determining tandem repeat organization in pericentric heterochromatin. These assemblies provide a platform for evolutionary and functional genomic studies of satDNA.

## MATERIALS AND METHODS

The detailed protocols for all molecular methods are available on our website (http://blogs.rochester.edu/larracuente/lab-protocols) and all computational pipelines and intermediate files are available on our lab Github page (https://github.com/LarracuenteLab/Khost_Eickbush_Larracuente2016).

### Assemblies

We downloaded raw and error-corrected SMRT PacBio sequence reads from the ISO1 strain of *D. melanogaster* (Kim, et al. 2014; raw read SRA accession SRX499318). We downloaded two assemblies constructed using the PBcR pipeline: *1)* “PBcR-BLASR”—an assembly made using Celera 8.1 and a computationally intensive all-by-all alignment with BLASR (Sergey Koren and Adam Phillipy); and *2)* “PBcR-MHAP”—an assembly made using Celera 8.2 and the minhash alignment process (MHAP; (Berlin, et al. 2015)).

We generated new assemblies using the PBcR pipeline from Celera 8.2 and 8.3 (MHAP) to explore the parameter space that produces the best assembly of repetitive loci (Table 1; Table S1). We tested 39 combinations of k-mer size, sketch size, and coverage, each with and without the large/diploid genome parameters (http://wgsassembler.sourceforge.net/wiki/index.php/PBcR#Assembly_of_Corrected_Sequences) that allows a more permissive error rate (Supplementary Files 1 and 2). We also created assemblies using the recently developed Canu 1.2 pipeline (Supplementary File 3). As Canu also uses the MHAP algorithm to overlap reads similar to the Celera 8.2+ pipeline, we attempted to use parameter settings that we had optimized for MHAP (kmer=14, sensitivity=high). We used a range of values for the master errorRate parameter, which implicitly sets other error rates (Table S3). In addition to the Celera assembler, we tested different parameter combinations in the experimental diploid PacBio assembler Falcon (https://github.com/PacificBiosciences/FALCON). We tested a range of -min_cov lengths, which controls the minimum coverage when overlapping reads in the pre-assembly error correction step, and a range of -min_len sizes, which sets the minimum length of a read to be used in assembly. Overall, we tested 19 different combinations (example spec files in Supplementary File 4 here: https://github.com/LarracuenteLab/Khost_Eickbush_Larracuente2016). All combinations of FALCON parameters produced a highly fragmented *Rsp* locus (Table S2), and thus were excluded from further analysis. We ran all assemblies on a node with a pair of Intel Xeon E5-2695 v2 processors (24 cores) and 124 GB on a Linux supercomputing cluster (Center for Integrated Research Computing, University of Rochester) using the SLURM job management system (http://slurm.schedmd.com/). Example specification files and SLURM scripts are found at https://github.com/LarracuenteLab/Khost_Eickbush_Larracuente2016 (*e.g.* Supplementary File 5). Not all parameter combinations resulted in finished assemblies, as numerous parameter combinations exceeded their allotted memory and failed, and others resulted in impractically long assembly times. For those that did finish, we evaluated assemblies for their ability to generate large contiguous blocks of the *Rsp* and *260-bp* satellites.

To determine the step in the assembly process that leads to the most contiguous assembly of repeats, we assembled reads corrected with the Celera 8.1 pipeline by BLASR (from Adam Phillipy and Sergey Koren; http://bergmanlab.ls.manchester.ac.uk/?p=2151) using the MHAP algorithm implemented in Celera 8.3 and the Canu 1.2 pipelines (Supplementary Files 6 and 7). In each case, we sampled the longest 25X subset of the BLASR-corrected reads, which we then converted to an .frg file and assembled using Celera 8.3 (“BLASR-corr Cel8.3”) or Canu 1.2 (“Canu-corr”).

### Assembly evaluation

We used custom repeat libraries that we compiled from Repbase (Supplementary File 8) and updated with consensus sequences of *1.688* family and *Responder* (*Rsp*) satellites as BLAST (blast/2.2.29+) queries against all assemblies. We created a custom Perl script to annotate contigs containing repetitive elements based on the BLAST output. The gff files containing our repeat annotations for the PBcR-BLASR assembly are in Supplementary Files 9-11 here: https://github.com/LarracuenteLab/Khost_Eickbush_Larracuente2016. For *Rsp*, we categorized repeats as either *Left, Right, Variant*, or *Truncated* based on their length and BLAST score. Our cutoff value to categorize *Rsp* repeats as *Left* or *Right* corresponds to the 90th percentile of the BLAST score distribution in reciprocal BLAST searches. We categorized *Rsp* repeats with a score below this cutoff as *Variant* and partial repeats <90 bp as *Truncated*. We evaluated PacBio assemblies based on the copy number and contiguity of *Rsp* and *260-bp* repeats (Table 1; Table S1). For both the *Rsp* and *260-bp* loci, we imported our custom gff files into the Geneious genome analysis tool (http://www.geneious.com; Kearse, et al. 2012) and manually annotated repeats that were still ambiguous. We also compared these assemblies to the *D. melanogaster* reference genome v6.03 (Hoskins, et al. 2015).

Cytological validation: We confirmed the higher-order genomic organization of *Rsp* and *260-bp* with fluorescence *in situ* hybridization (FISH). We designed a Cy5-labeled oligo probe to the *Bari1* repeats distal to the *Rsp* locus (*Bari1:* 5’-/Cy-5/ATGGTTGTTTAAGATAAGAAGGTATCCGTTCTGAT-3’) (Fig S1). We generated biotin- and digoxigenin-labeled probes using nick translation on gel-extracted PCR products from the *Rsp* and *260-bp* repeats, respectively (260F: 5’-TGGAAATTTAATTACGAGCT-3’; 260R: 5’-ATGAAACTGTGTTCAACAAT-3’; (Abad, et al. 2000); RspF: 5’-CCGATTTCAAGTACCAGAC-3’; RspR: 5’-GGAAAATCACCCATTTTGACCGC-3’; (Larracuente 2014). We conducted FISH according to (Larracuente and Ferree 2015; Fig 1; Fig S1). Briefly, larval brains were dissected in 1X PBS, treated with a hypotonic solution (0.5% Sodium citrate) and fixed in 1.8% paraformaldehyde; 45% acetic acid and dehydrated in ethanol. Probes were hybridized overnight at 30°C, washed in 4X SSCT and 0.1X SSC, blocked in a BSA solution and treated with 1:100 Rhodamine-avadin (Roche) and 1:100 anti-dig fluorescein (Roche), with final washes in 4X SSCT and 0.1X SSC. Slides were mounted in VectaShield with DAPI (Vector Laboratories), visualized on a Leica DM5500 upright fluorescence microscope at 100X, imaged with a Hamamatsu Orca R2 CCD camera and analyzed using Leica’s LAX software.

Computational validation: Because we only use a subset of error-corrected PacBio reads to create *de novo* assemblies, we assessed the computational support for each assembly using independently derived short Illumina reads, Sanger-sequenced BACs and the entire set of raw PacBio reads. We mapped high-coverage Illumina reads from the ISO1 strain (Gutzwiller, et al. 2015) to each assembly using “–very-sensitive” settings in bowtie2 (Langmead and Salzberg 2012) to identify regions of low coverage that could indicate mis-assemblies (*eg.* Figs S4 and S5). We quantified error rate with these Illumina reads using Pilon 1.2 (Walker, et al. 2014) and Bcftools 0.1.19. We calculated the number of nucleotide substitutions/contig length and the number of indels/contig length for each contig in the assembly and plotted the distribution (Figs. S10 and S11). We mapped raw PacBio reads to the PBcR-BLASR, BLASR-corr Cel 8.3, MHAP 20_25X, and Canu 4% error rate assemblies using the default parameters in the PacBio-specific BLASR aligner in the SMRT Analysis 2.3 software package available from Pacific Biosciences (*eg.* Fig S7 and S8). For the BLASR-corr Cel 8.3, MHAP 20_25X, and Canu 4% error rate assemblies, we provided the mapped PacBio reads to the Quiver genomic consensus caller to correct remaining SNPs/indels (https://github.com/PacificBiosciences/GenomicConsensus). We also mapped to our assemblies available BACs sequences (BACN05C06, BACR32B23, CH221-04O17) that localize to the *Rsp* locus (Larracuente 2014).

Molecular validation: We confirmed the presence of two distinct G5 clusters using PCR analysis with primers designed in and around informative indels (Fig 2A). To confirm the locus orientation, we designed PCR primers that could only amplify an ~15 kb segment of the distal part of the locus found in the PBcR-BLASR assembly (Fig 2A; primer pair 3; Fig S2A) and confirmed its organization using restriction enzyme digestion with *HindIII*, *EagI*, *SstI*, and *XmaI* and Southern blot analysis using a biotinylated *Rsp* probe and the North2South kit (ThermoFisher #17175, Fig S2B). We validated the distal and proximal ends of the locus with a Southern blot analysis (see below) on genomic DNA digested with *AccI, EcoRI, FspI* and *SstI*.

### Composition and structure of satellite loci

Using maps of the locus based on our BLAST output, we extracted individual repeat units and created alignments using ClustalW (Larkin, et al. 2007). We inspected and adjusted each alignment by hand in Geneious 8.05 (Kearse, et al. 2012) and examined the relationship between genetic distance and physical distance between repeats. We used the APE phylogenetics package in R (Paradis, et al. 2004) to construct neighbor-joining trees for all monomers of each repeat family, using the “indelblock” model of substitution (Fig 3). We then collapsed the repeats down to individual unique variants and plotted their distribution across the locus using a custom Perl script to examine any higher-order structures (Fig 4). All scripts are available at https://github.com/LarracuenteLab/Khost_Eickbush_Larracuente2016.

### Southern Blot Analyses

Spooled genomic DNA was obtained from ~60 adult females in standard phenol-chloroform extractions and re-suspended in TE buffer. We performed Southern blot analyses on ~10 ng of the 15-kb PCR amplicon and 10 μg of genomic DNA. In short, restriction enzyme digested DNA was fractionated on a 1% agarose/TAE gel and then depurinated, denatured, and neutralized before being transferred for 16 hrs in high salt (20 X SSC/1 M NH4Acetate) to a nylon membrane (Genescreen PlusR). DNA was UV crosslinked and hybridizations were done overnight at 55°C in North2South hybridization buffer (ThermoScientific). To make the biotinylated RNA probe, we transcribed a 240-bp *Rsp* gel extracted PCR amplicon (primers: T7_rsp1 5’-TAATACGACTCACTATAGGGGAAAATCACCCATTTTGATCGC-3” and rsp2 5’-CCGAATTCAAGTACCAGAC-3’) using the Biotin RNA Labeling Mix (Roche) and T7 polymerase (Promega). The hybridized membrane was processed as recommended for the Chemiluminescent Nucleic Acid Detection Module (ThermoScientific), and the signal recorded on a ChemiDoc XR+ (BioRad). For the slot blot protocol, see Supplemental Methods.

### Nuclei Isolation and Pulse-Field Gel Analysis

Nuclei isolation was performed as described in (Kuhn, et al. 2008) with some modification. Approximately 100 flies were ground in liquid nitrogen. The powder was suspended in 900 μl of nuclei isolation buffer with 5 mM DTT, filtered first through a 50-μm and then through a 20-μm nitex nylon membrane (03-50/31 and 03-20/14, Sefar America) and pelleted by centrifugation at 3500 rpm for 10 mins. Nuclei were re-suspended in 200 μl of 30 mM Tris, pH 8.0, 100 mM NaCl, 50 mM EDTA, 0.5% Triton X-100; combined with an equal volume of 1% agarose and set using a block maker (BioRad). The agarose blocks were incubated in 0.5 M EDTA (pH 8.0), 1% sodium lauryl sarcosine, 0.1 mg/ml proteinase K overnight at 50°C and then washed in TE and restriction enzyme buffer. The blocks were digested overnight in fresh buffer with BSA and 100 units of *EcoRI* and *AccI* at 37°C. The digested blocks were run in a 1% agarose/TBE gel using a pulse field apparatus for 21 hrs at 8°C (4.5 V/cm; 0.5-50 sec pulses). Southern analysis was performed as above using the biotinylated *Rsp* probe.

## ACKNOWLEDGMENTS

We would like to thank Casey Bergman for helpful conversations about PacBio assembly methods and for sharing assemblies, reads and protocols. We would like the thank the staff of the Center for Integrated Research Computing at the University of Rochester for maintenance of the computing cluster and access to computational resources and Tom Eickbush for discussion. This work was supported by the University of Rochester.

## Supplemental methods

### Slot Blots

Genomic DNA (100 ng to 600 ng) was denatured (final concentration 0.25 N NaOH, 0.5 M NaCl) for 10 mins at room temperature and then quick cooled. Slot blots were performed as recommended using a 48-well BioDot SF microfiltration apparatus (Bio-Rad). Each blot was first hybridized with a biotinylated rp49 RNA probe (generated as per above with primers: T7_rp49REV 5’-GTAATACGACTCACTATAGGGCAGTAAACGCGGTTCTGCATG-3 and rp49FOR 5’-CAGCATACAGGCCCAAGATC-3’). The membrane was then stripped with a 100° C solution of 0.1X SSC/0.5% SDS (3 times for ~20 mins) and re-hybridized with the *Rsp* probe as per the above Southern analysis. Signals were quantitated using the ImageLab software (BioRad).

**Figure S1.**
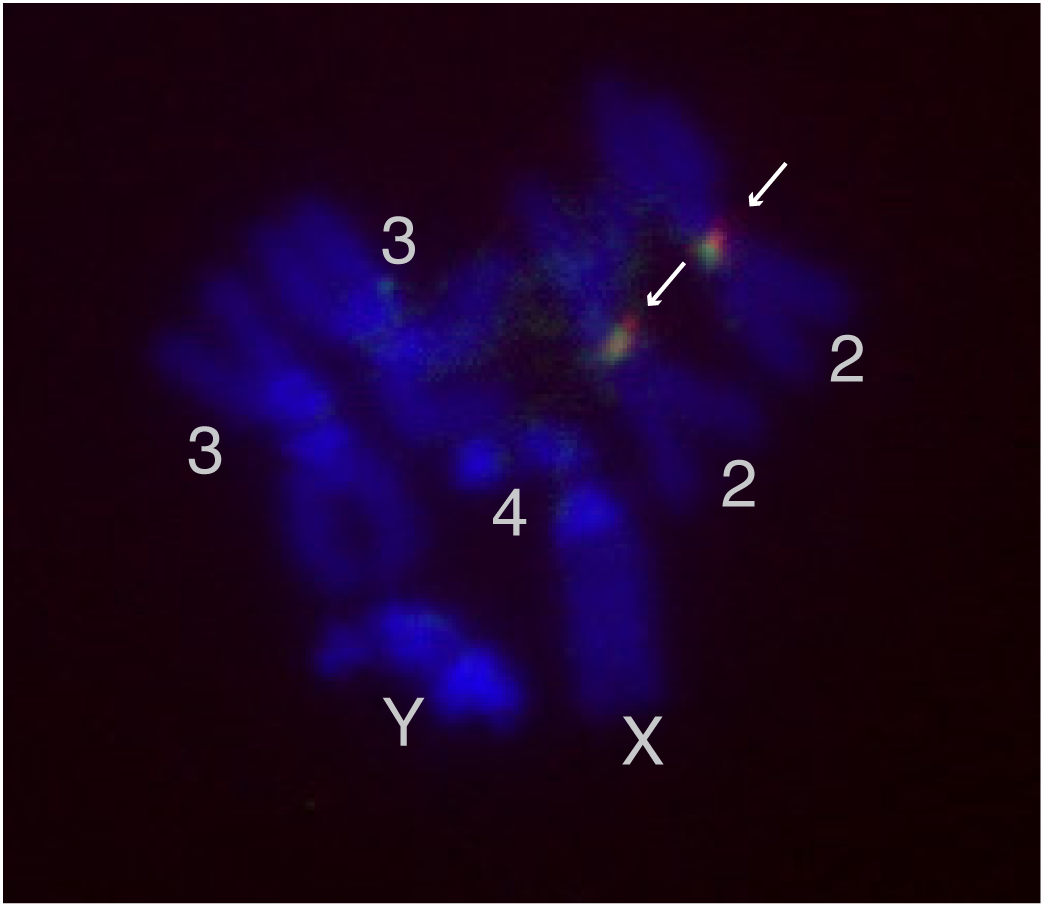
*Rsp* and *Bari1* clusters on chromosome *2R*. Fig S1: FISH image of *D. melanogaster* mitotic chromosomes from 3^rd^ instar larvae. DNA is stained with DAPI (blue). *Bari-1* repeats are labeled with an anti-digoxigenin probe (green), which co-localizes closely with *Rsp* repeats labeled with an avidin-rhodamine probe (red).

**Figure S2:**
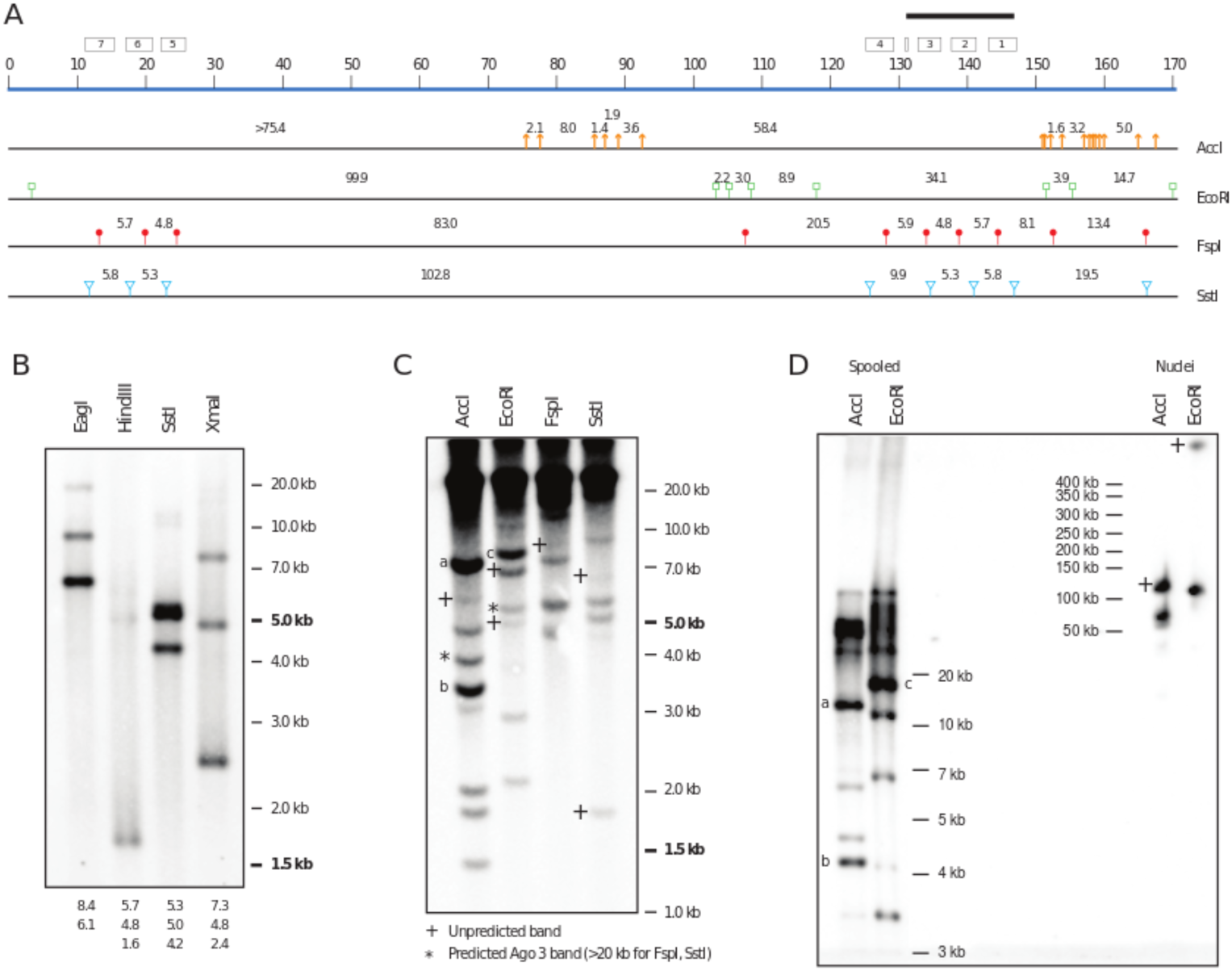
Molecular validation of *Rsp* locus. Figure S2: A. Schematic showing the extent of the assembled *Rsp* array (blue line as in Fig 2A) locations of *AccI, EcoRI, FspI*, and *SstI* sites within the array. Predicted fragment sizes based on *in silico* digestion of the *Rsp* locus in the PBcR-BLASR are also indicated. Because there are additional *Rsp* repeats proximal (left on the diagram, see text) to the assembled array, it is not possible to determine the size of the proximal fragment(s) for any of the restriction enzymes (*e.g.* a proximal *AccI* fragment is >75 kb). The position of *G5 Jockey* repeats are indicated by boxes. The extent of the 15-kb long PCR fragment used in (B) is indicated by a black bar. B. The 15-kb PCR product was digested with the indicated enzymes, and the separated fragments blotted and hybridized with a *Rsp* probe. The sizes of predicted fragments (with *Rsp* repeats) generated as described in (A) are listed below the blot. C. Genomic DNA was digested, and the separated fragments blotted and hybridized with a *Rsp* probe. Only fragments < 20 kb in size can be resolved. Band sizes can be compared to the *in silico* predictions in (A). Expected bands from the *Ago3* locus on *3L* are indicated by an asterisk. Unpredicted fragments are indicated with a + and could represent hybridization to fragments proximal to the array. Select bands are indicated with a letter (a, b,c) and their sizes compared in (D). D. Pulse-field gel of spooled DNA (left) and high molecular weight DNA (nuclei, right) digested and probed with *Rsp*. The intensity of the bands labeled a, b, c (left) anchor the patterns seen in (C) but the *Rsp* fragments appear to run slower under the pulse field conditions. For right lanes, the size of the smaller fragment in each digest can be compared to the *in silico* prediction. + as in (C).

**Fig. S3:**
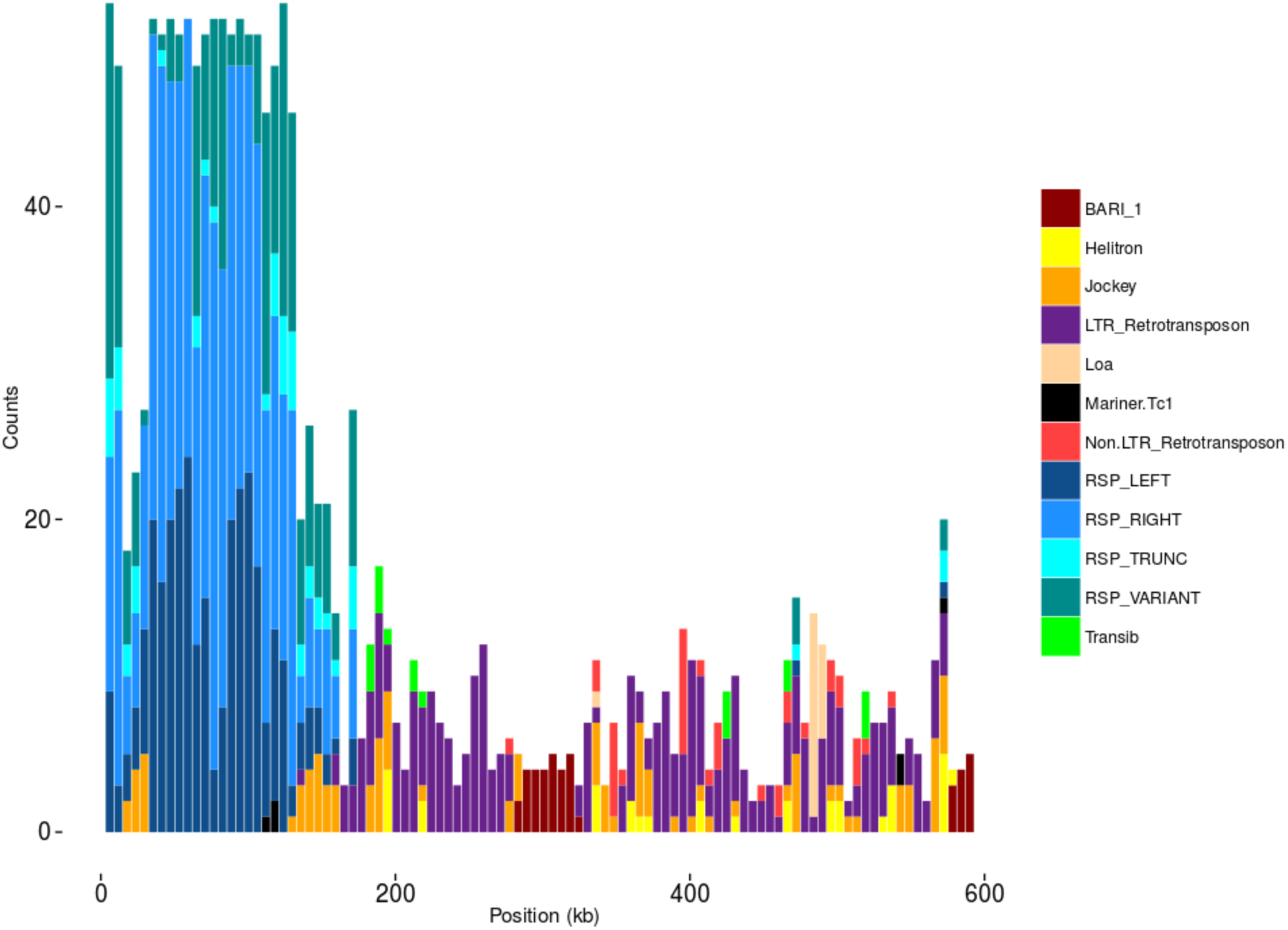
Major *Rsp* locus in Canu 4% assembly. Figure S3: Major *Rsp* locus in Canu 4% assembly. Counts for each element in our custom Repbase library in 5-kb windows across the *Rsp* locus in our Canu 4% assembly. The main block of *Rsp* repeats is approximately equal in repeat number to the PBcR-BLASR assembly, and orientation of the repeats is supported by restriction digest and Southern blot analysis. The contig extends the PBcR-BLASR contig ~300 kb distally, which includes an additional 10 variant *Rsp* repeats (minor *Rsp* locus) as well as the *Bari1* clusters. These additional repeats are also present in the PBcR-BLASR assembly, but on a separate contig.

**Figure S4:**
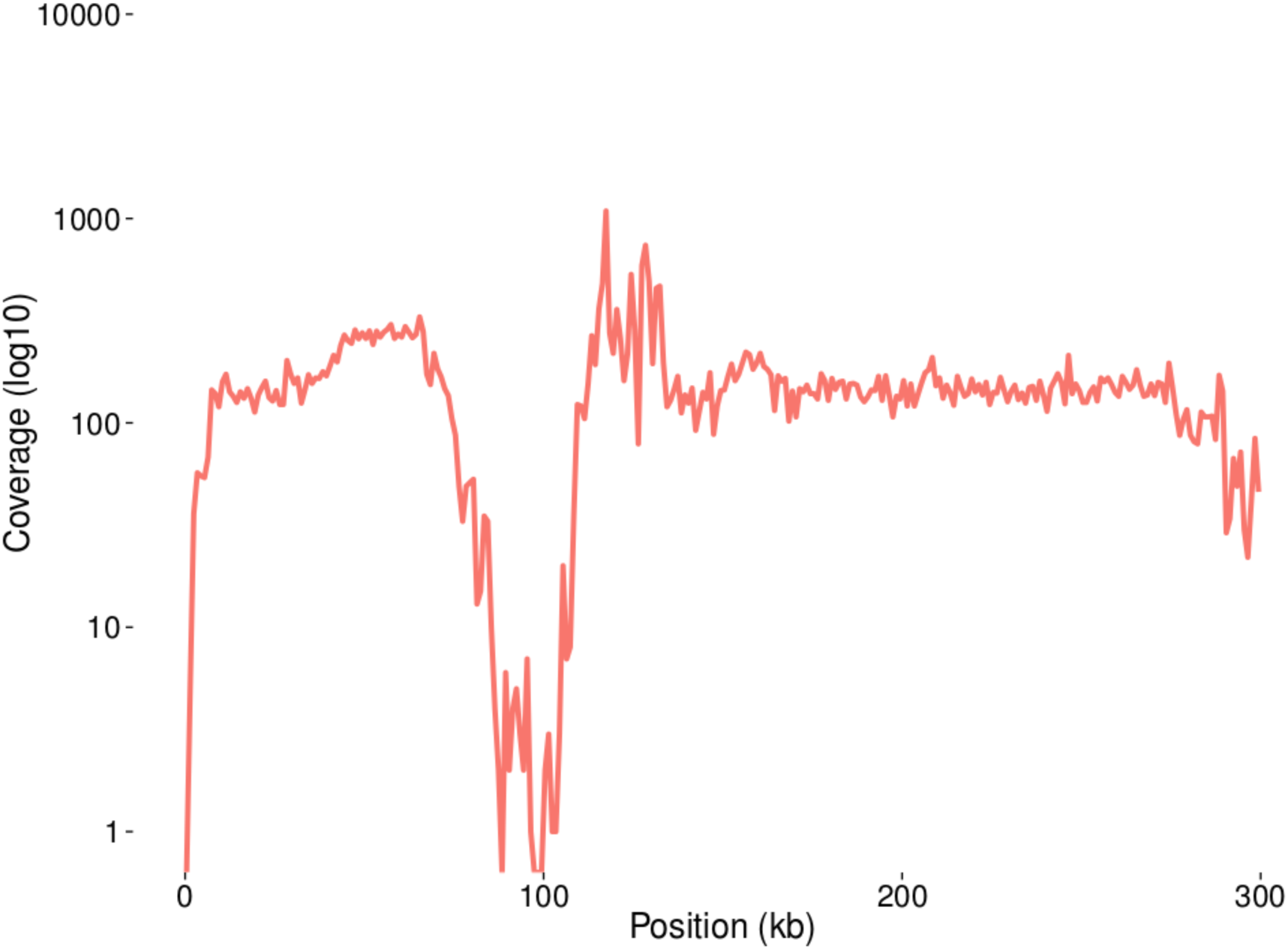
Coverage of deep Illumina reads over *Rsp* locus in the MHAP 20_1500_25X assembly. Figure S4: Coverage of deep Illumina reads (SRA accession ERR701706) over *Rsp* locus in MHAP 20_1500_25X assembly mapped using the –very-sensitive settings in Bowtie2. Coverage was plotted in 1-kb windows using Bedtools and is shown on a log scale.

**Fig. S5:**
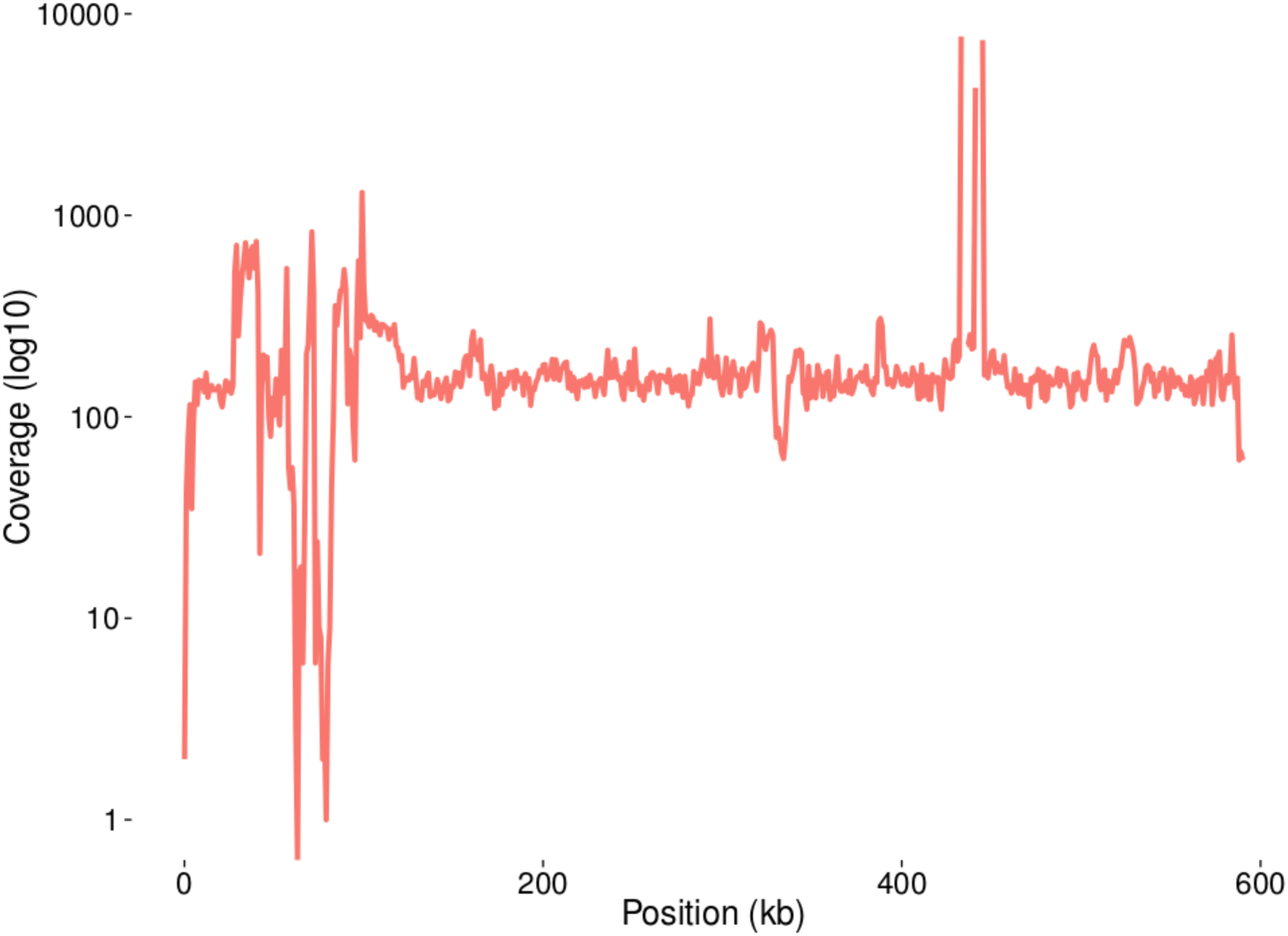
Coverage of deep Illumina reads over *Rsp* contig in Canu 4% assembly. Figure S5: Coverage of deep Illumina reads (SRA accession ERR701706) over *Rsp* locus in Canu 4% error-rate assembly mapped using –very-sensitive settings in Bowtie2. Coverage was plotted in 1-kb windows using Bedtools and is shown on a log scale.

**Figure S6:**
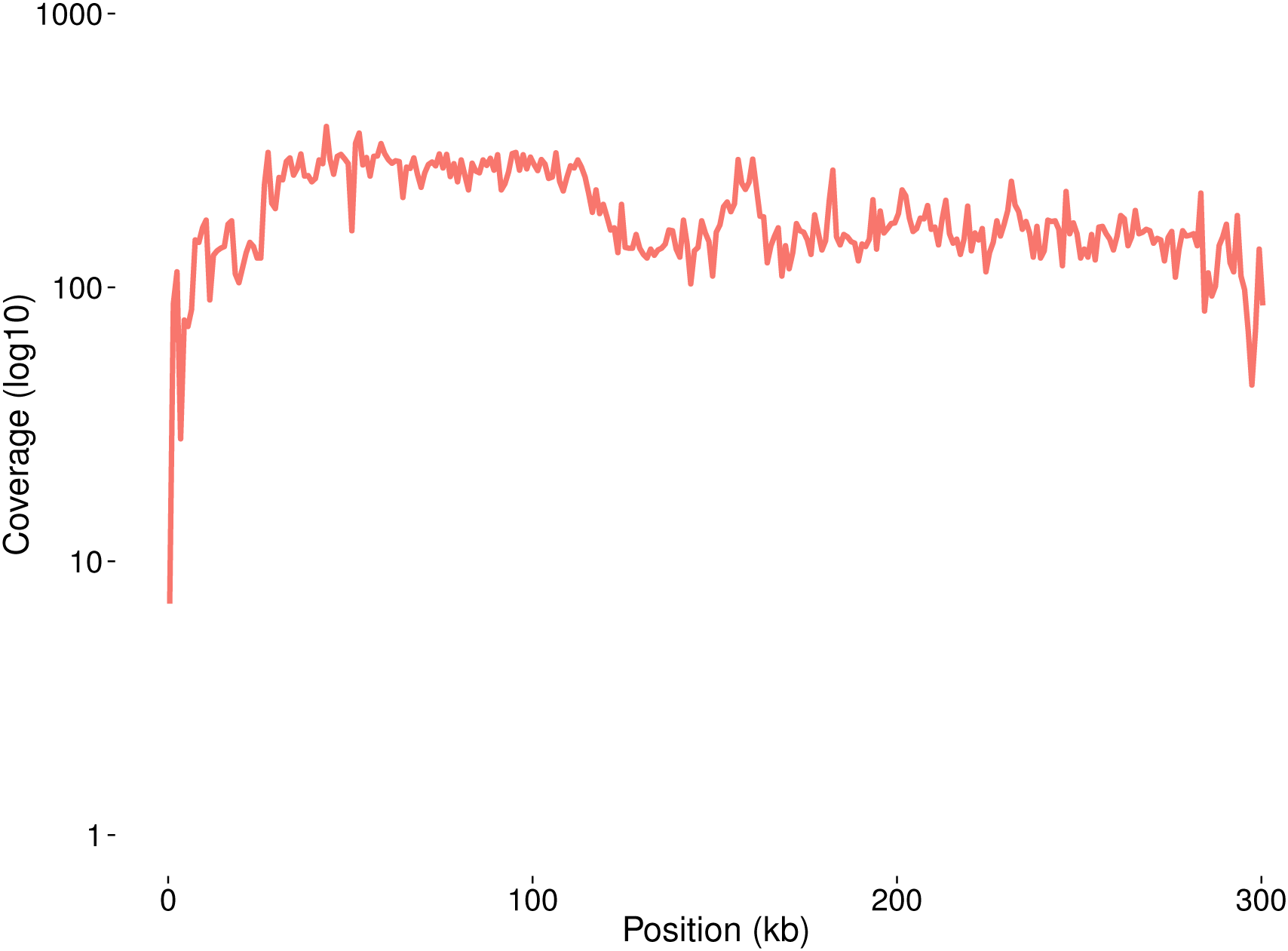
Coverage of deep Illumina reads over *Rsp* locus in PBcR-BLASR assembly. Figure S6: Coverage of deep Illumina reads (SRA accession ERR701706) over *Rsp* locus in PBcR-BLASR assembly mapped using the –very-sensitive settings in Bowtie2. Coverage was plotted in 1-kb windows using Bedtools and is shown on a log scale.

**Figure S7:**
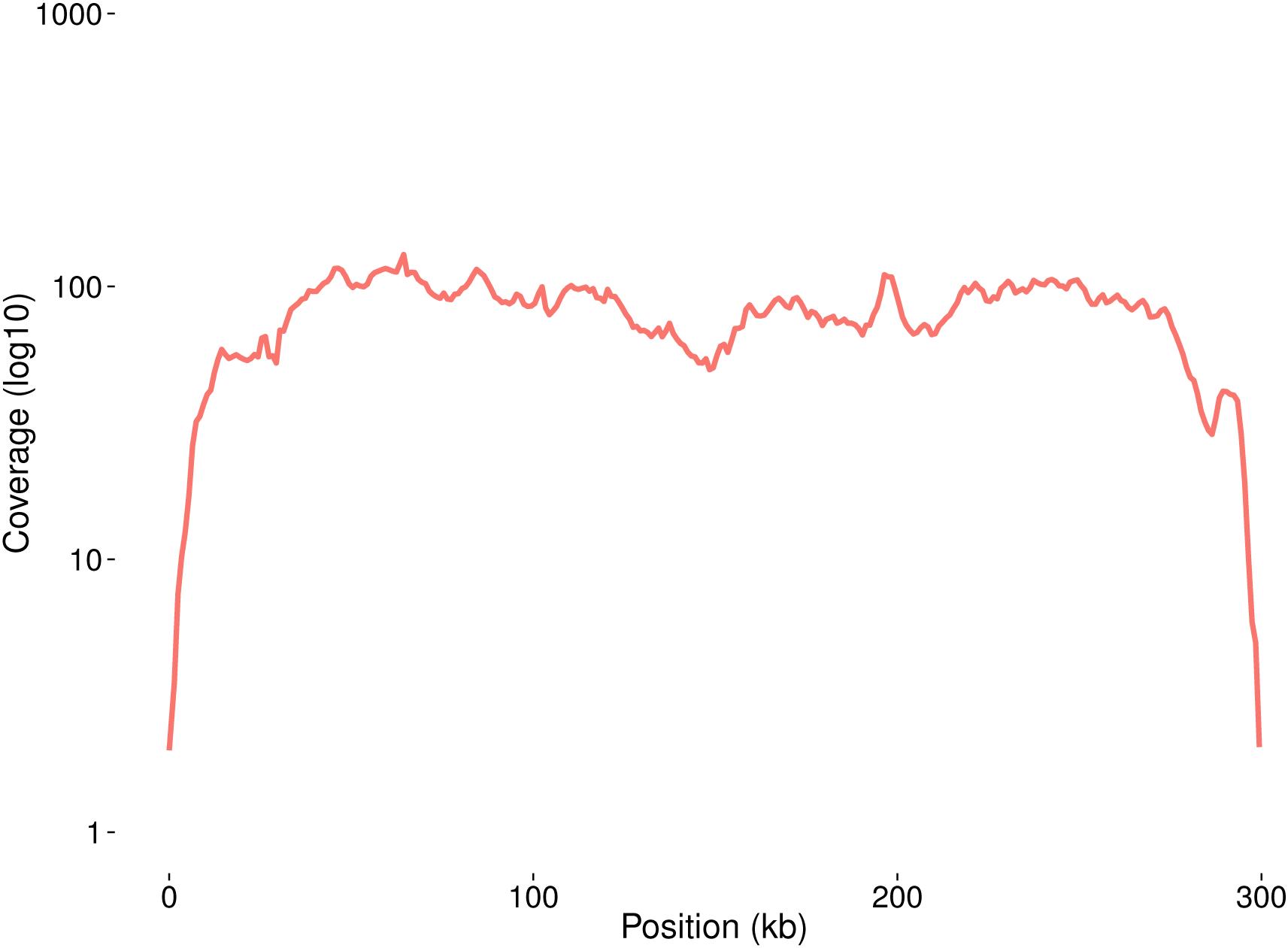
Coverage of raw PacBio reads mapped against PBcR-BLASR assembly. Figure S7: Coverage of raw PacBio reads mapped against PBcR-BLASR assembly. Reads were mapped using BLASR included in SMRT Analysis v. 2.3. Coverage shown on a log(10) scale.

**Figure S8:**
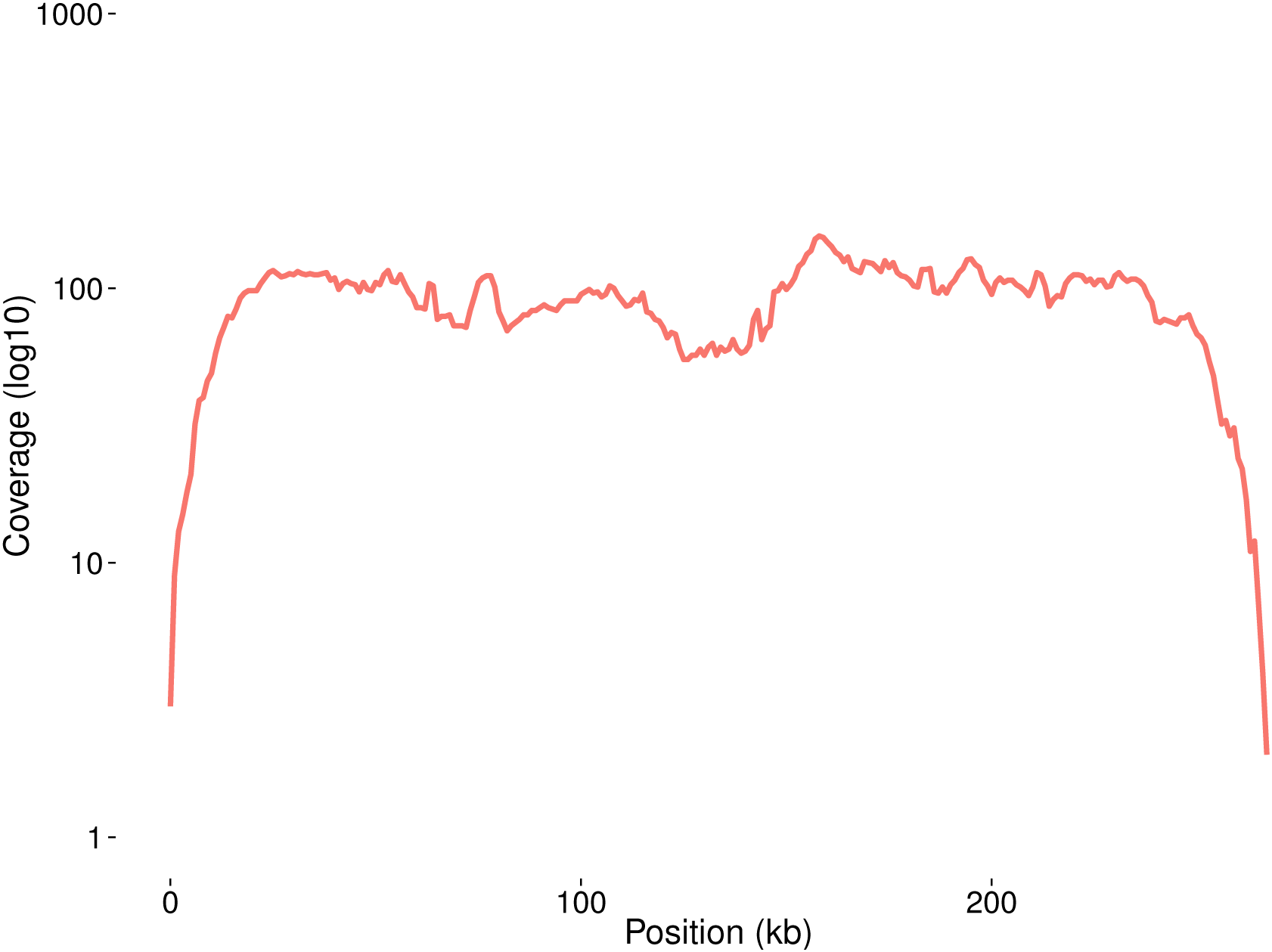
Coverage of raw PacBio reads mapped against BLASR-corr Cel8.3 assembly. Figure S8: Coverage of raw PacBio reads mapped against BLASR-corr Cel8.3 assembly. Reads were mapped using BLASR included in SMRT Analysis v. 2.3. Coverage shown on a log(10) scale.

**Figure S9:**
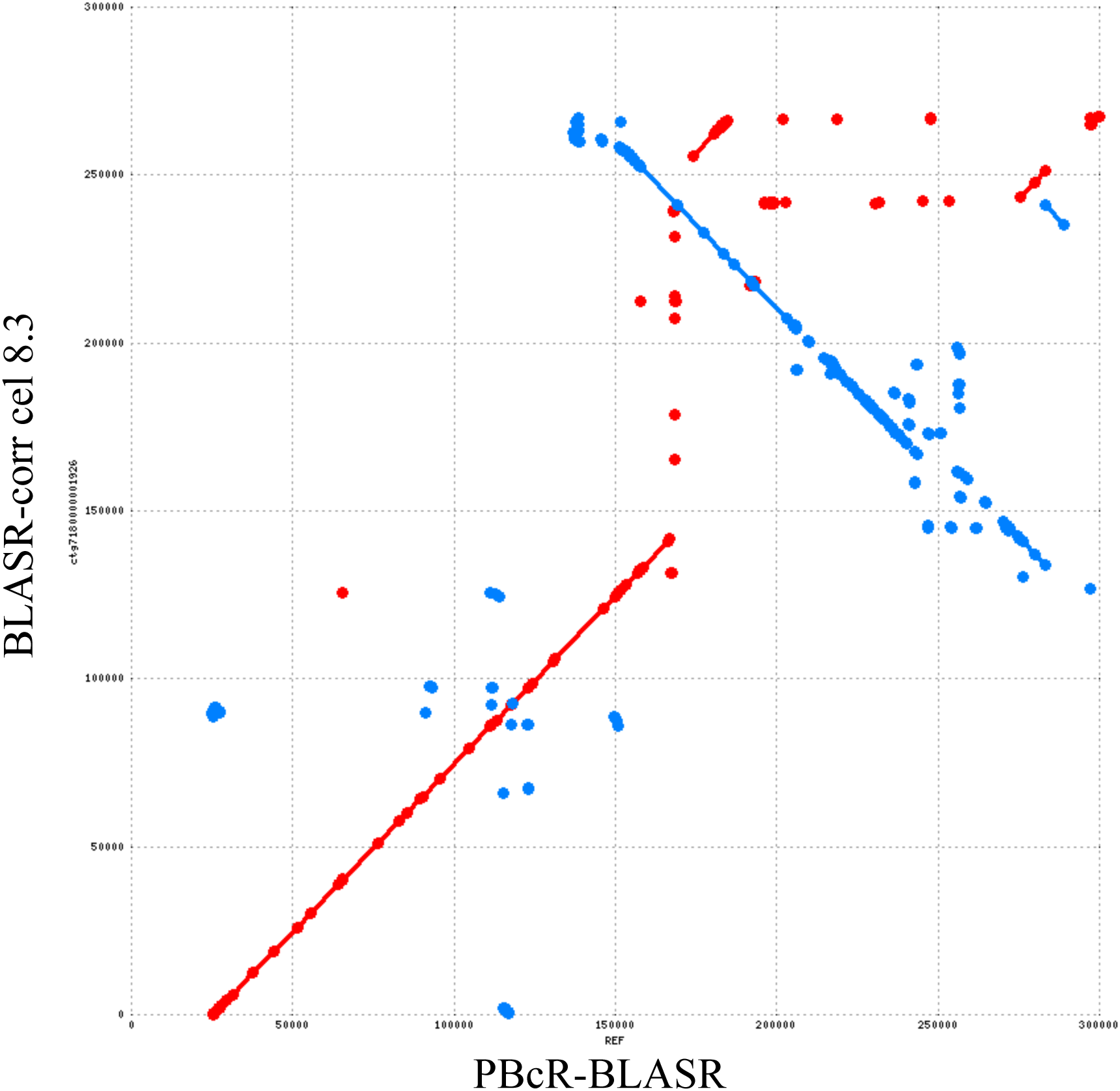
Mummer dotplot alignment of *Rsp* loci. Figure S9: Mummer dotplot alignment of *Rsp* loci contigs from the two best-supported assemblies (PBcR-BLASR and BLASR-corr Cel8.3). Red points indicate identical strand, blue indicates opposite strand.

**Fig. S10:**
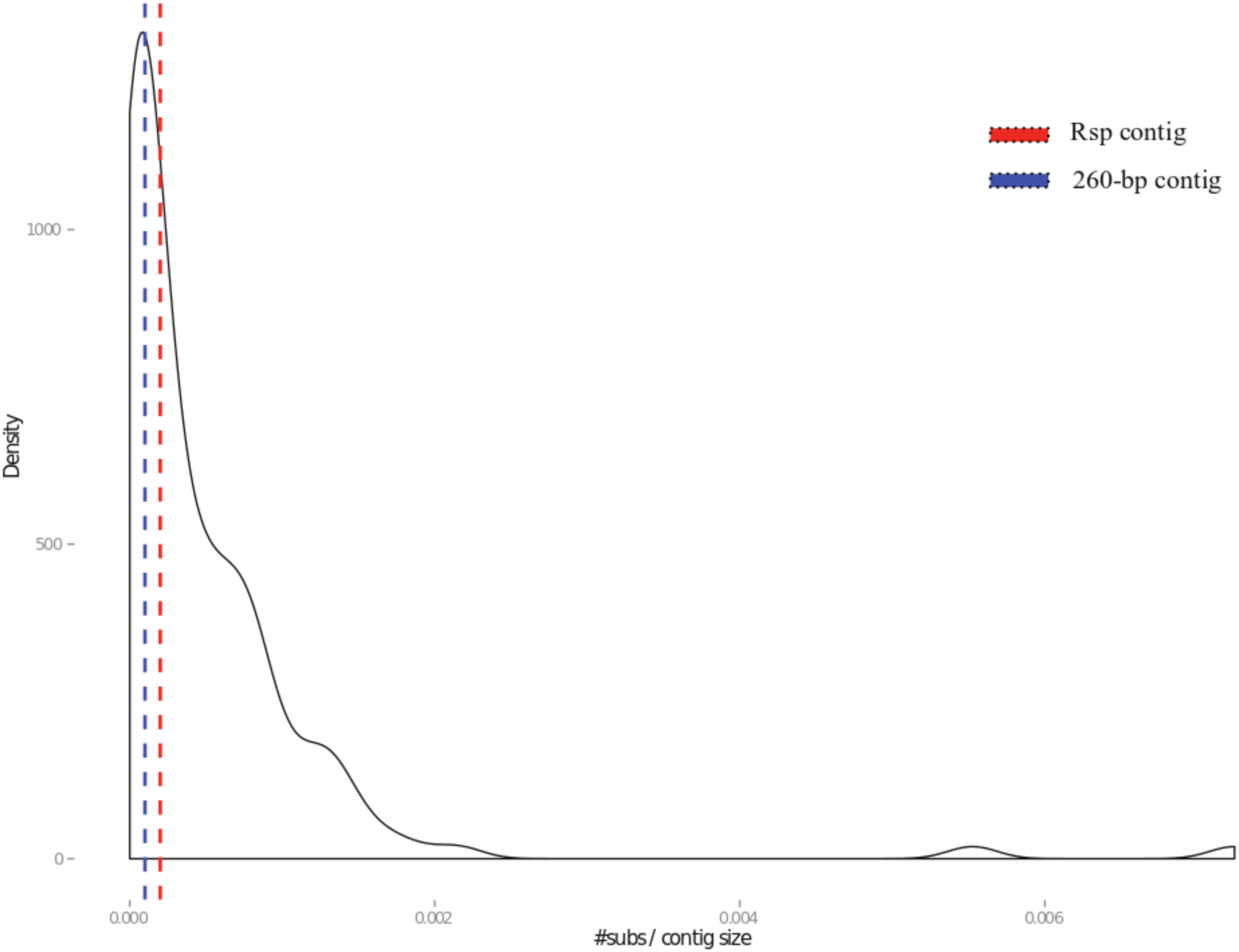
Nucleotide substitution error distribution. Figure S10: Density plot showing distribution of nucleotide substitutions fraction for each contig in the PBcR-BLASR assembly. Illumina reads were mapped to the assembly using Bowtie2, and Pilon was used to generate variant call format (vcf) files. SNPs for each contig were summed and divided by the total contig length.

**Fig S11:**
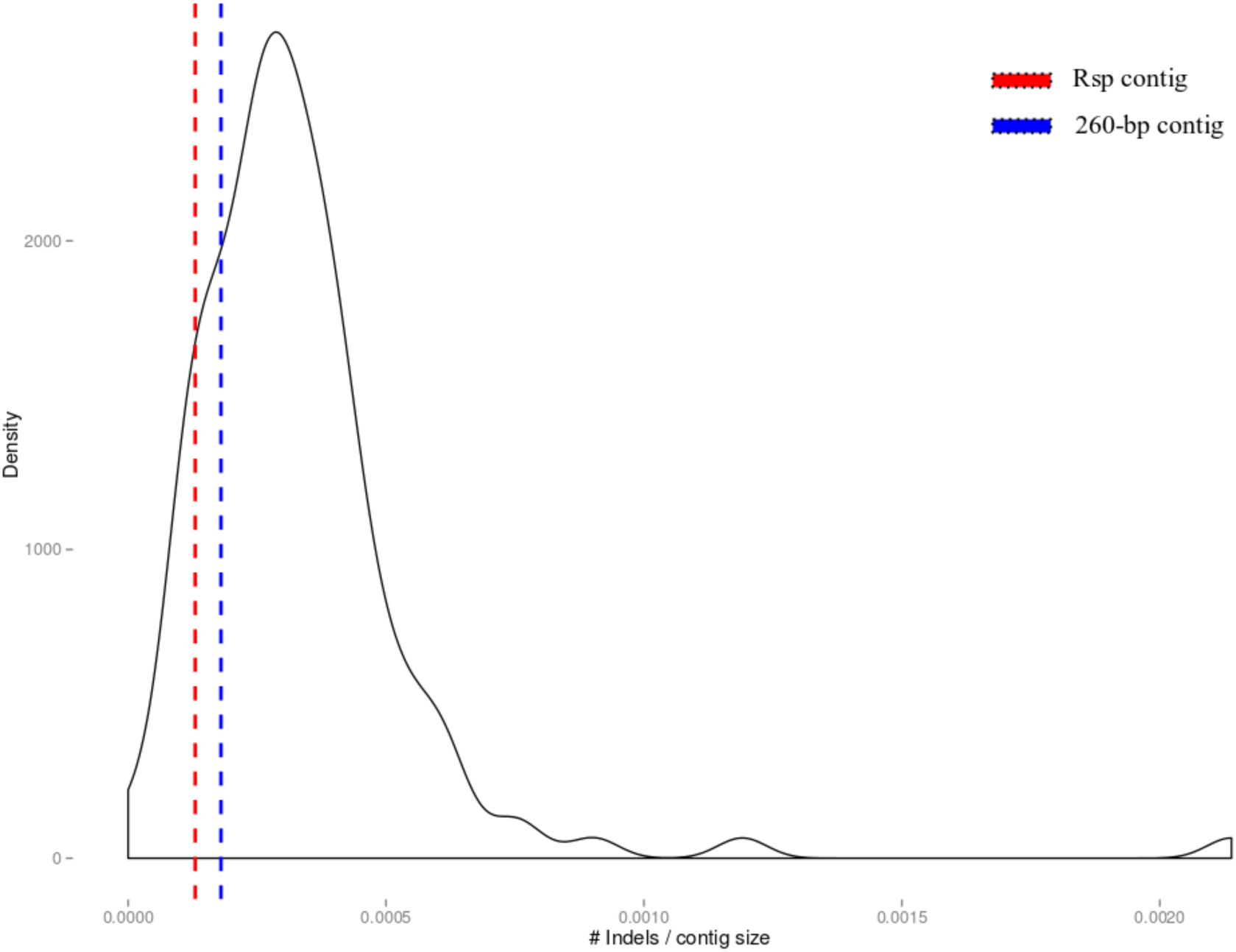
Indel distribution. Figure S11: Density plot showing distribution of indel fraction for each contig in the PBcR-BLASR assembly. Illumina reads were mapped to the assembly using Bowtie2, and Pilon was used to generate variant call format (vcf) files. Indels for each contig were summed and divided by the total contig length.

**Figure S12:**
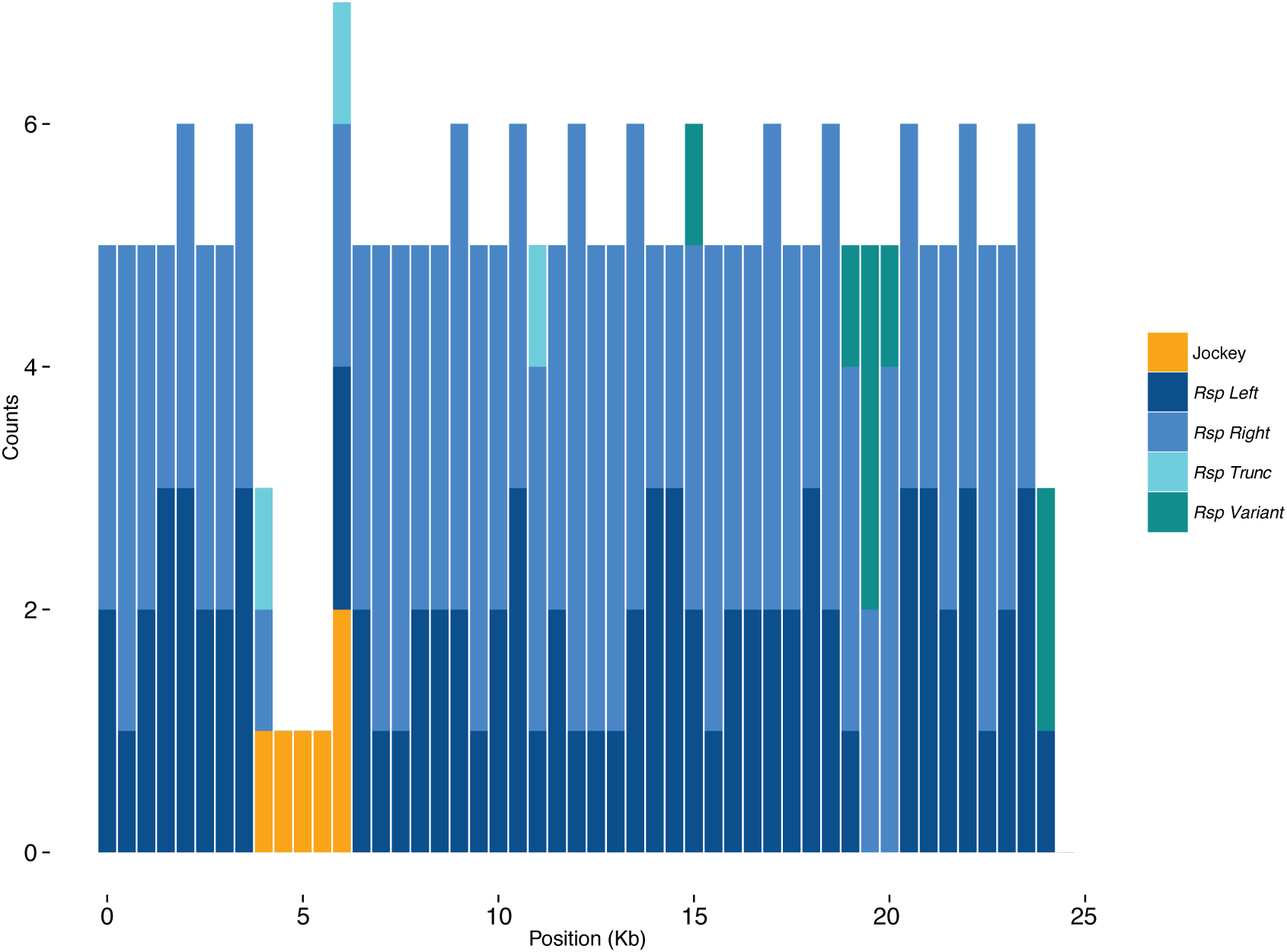
Map of centromere-proximal *G2* contig. Figure S12: Map of the centromere-proximal *G2-Rsp* contig assembled using corrected PacBio reads. Counts for each repetitive elements in our custom Repbase library are plotted in 500 bp windows across the contig.

**Fig S13:**
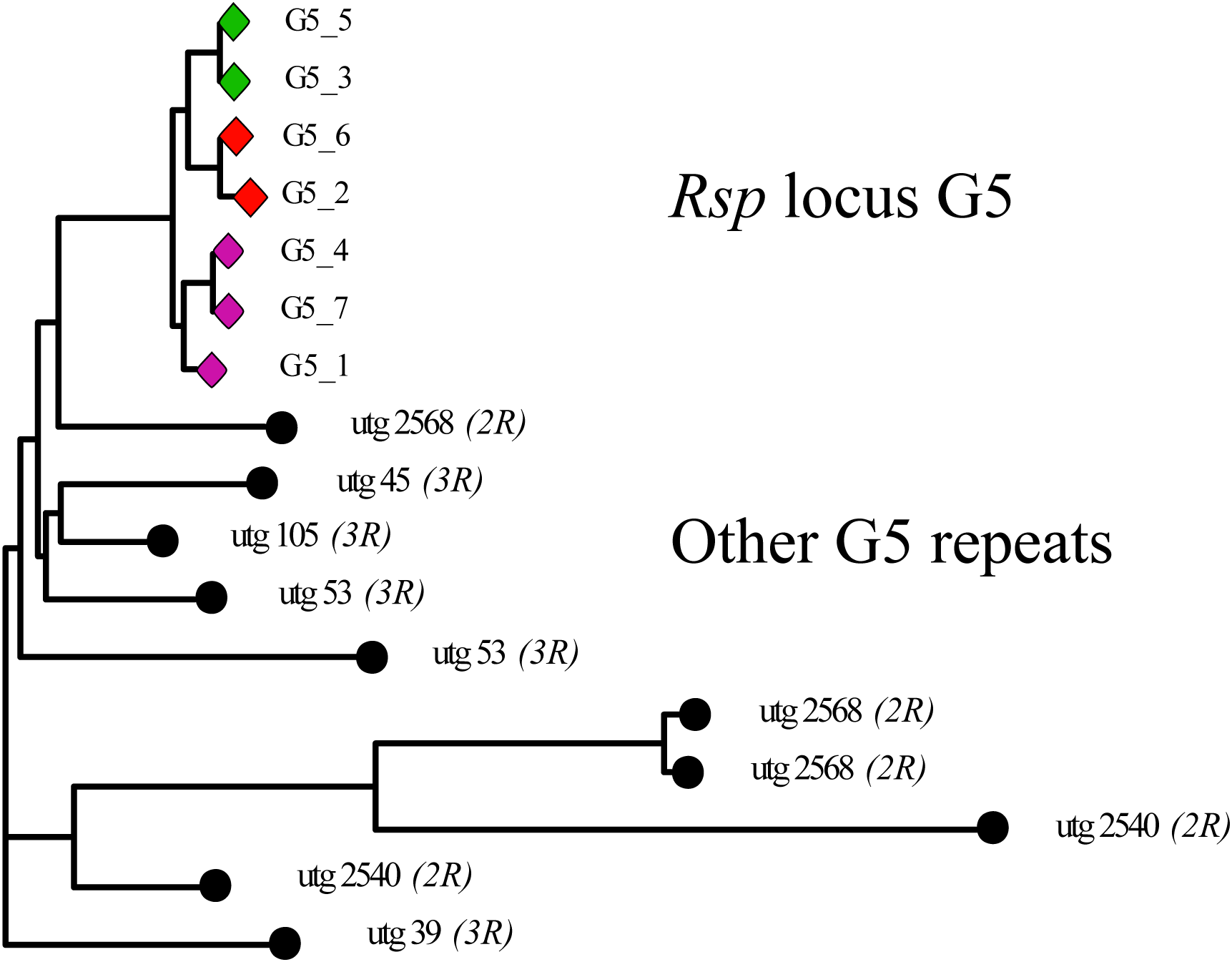
G5 phylogeny. Figure S13: RaxML tree of G5 repeats at *Rsp* locus and all other *G5* repeats > 1000bp in the PBcR-BLASR assembly. Tree was built in Geneious with 100 bootstraps. The other *G5* repeats on *2R* are all at least 2 Mb distal to the *Rsp* locus *G5* cluster.

